# Continuous Evaluation of Denoising Strategies in Resting-State fMRI Connectivity Using fMRIPrep and Nilearn

**DOI:** 10.1101/2023.04.18.537240

**Authors:** Hao-Ting Wang, Steven L Meisler, Hanad Sharmarke, Natasha Clarke, Nicolas Gensollen, Christopher J Markiewicz, François Paugam, Bertrand Thirion, Pierre Bellec

## Abstract

Reducing contributions from non-neuronal sources is a crucial step in functional magnetic resonance imaging (fMRI) connectivity analyses. Many viable strategies for denoising fMRI are used in the literature, and practitioners rely on denoising benchmarks for guidance in the selection of an appropriate choice for their study. However, fMRI denoising software is an ever-evolving field, and the benchmarks can quickly become obsolete as the techniques or implementations change. In this work, we present a denoising benchmark featuring a range of denoising strategies, datasets and evaluation metrics for connectivity analyses, based on the popular fMRIprep software. The benchmark is implemented in a fully reproducible framework, where the provided research objects enable readers to reproduce or modify core computations, as well as the figures of the article using the Jupyter Book project and the Neurolibre reproducible preprint server (https://neurolibre.org/). We demonstrate how such a reproducible benchmark can be used for continuous evaluation of research software, by comparing two versions of the fMRIprep software package. The majority of benchmark results were consistent with prior literature. Scrubbing, a technique which excludes time points with excessive motion, combined with global signal regression, is generally effective at noise removal. Scrubbing however disrupts the continuous sampling of brain images and is incompatible with some statistical analyses, e.g. auto-regressive modeling. In this case, a simple strategy using motion parameters, average activity in select brain compartments, and global signal regression should be preferred. Importantly, we found that certain denoising strategies behave inconsistently across datasets and/or versions of fMRIPrep, or had a different behavior than in previously published benchmarks. This work will hopefully provide useful guidelines for the fMRIprep users community, and highlight the importance of continuous evaluation of research methods. Our reproducible benchmark infrastructure will facilitate such continuous evaluation in the future, and may also be applied broadly to different tools or even research fields.

## Introduction

Resting-state functional magnetic resonance imaging (fMRI) is a tool for studying human brain connectivity (Biswal et al., 2010; Fox & Greicius, 2010) which comes with many analytical challenges (Cole et al., 2010; Satterthwaite et al., 2012). One such key challenge is the effective correction of non-neuronal sources of fluctuations (called confounds), known as denoising, which is important to reduce bias when studying the association between connectomes and behavioral measures of interest (Chyzhyk et al., 2022). A wide range of denoising strategies have been proposed in the literature, with no approach emerging as a clear single best solution. Denoising benchmarks on functional connectivity (Ciric et al., 2017; Parkes et al., 2018) have thus become an important resource for the community to understand which denoising strategy is most appropriate in a given study. Denoising benchmarks are however at a constant risk of becoming obsolete, with new strategies being regularly developed or revised, as well as an ever-expanding scope of populations being enrolled in research studies. The main objective of the present work is to develop a fully reproducible denoising benchmark for fMRI functional connectivity, and demonstrate how the proposed infrastructure enables the continuous evaluation of denoising strategies across multiple software versions and datasets.

Reproducible and robust results have become a recurring interest in the neuroimaging community (Botvinik-Nezer et al., 2020; Niso et al., 2022). The popular package fMRIPrep (Esteban et al., 2019) is a prominent solution for fMRI preprocessing designed with reproducibility in mind, and we decided to build upon that software for our benchmark. However, fMRIPrep only performs minimal preprocessing while generating a broad list of potential confounds, intentionally leaving the selection of the exact denoising strategy to end-users. The connectivity metrics are also not included as part of fMRIPrep outputs, and users rely on additional software to apply denoising to time series and generate connectivity measures. One popular open-source Python library for this purpose is Nilearn (Abraham et al., 2014). Yet, until recently, there was no straightforward way to incorporate fMRIPrep outputs into Nilearn in order to reproduce the most common denoising strategies. This lack of integration represented a major barrier to the exploration of denoising tools, both for cognitive neuroscientists who were required to develop custom code, and for computer scientists who had to develop a detailed understanding of the inner workings of denoising strategies and fMRIPrep.

The main references for denoising benchmarks (Ciric et al. 2017, Parker et al., 2018) did not use the then-novel fMRIPrep. Whether the results of these benchmarks remain consistent with fMRIPrep outputs is an open question. Different fMRI preprocessing softwares provide largely similar results, but noticeable differences are still present (Bowring et al., 2019; Li et al., 2021). Other computational factors can possibly impact the conclusion of a benchmark, such as the version of software and operating system (Gronenschild et al., 2012). Recent research has also demonstrated that, given one fMRI dataset and similar research goals, different researchers will select a wide variety of possible analytical paths (Botvinik-Nezer et al., 2020). The lack of standard integration between fMRIPrep and Nilearn could lead to differences (and errors) in the implementation of the same denoising strategies by researchers, which can in turn lead to marked differences in the impact of denoising methods.

In this work, we propose to address the issues of robustness of functional connectivity denoising benchmarks by building a fully reproducible solution. This reproducible benchmark will allow the fMRI research community to consolidate past knowledge on technical advances, examine computation instability across different software versions, and provide guidance for practitioners. For the broader scientific research community, we aim to highlight the importance of continuous method evaluation, and propose a widely applicable infrastructure to implement such benchmarks. In order to create this benchmark, we implemented a series of specific objectives:

- First, we developed a standardized application programming interface (API) to extract nuisance regressors from fMRIPrep. The robust API, which was added to Nilearn release 0.9.0, can be used to flexibly retrieve a subset of fMRIPrep confounds for denoising and precisely replicate nuisance regressors based on denoising strategies proposed in the literature.
- Our second objective was to implement a denoising benchmark to provide recommendations on the choice of functional connectivity denoising strategies for fMRIPrep users. We used easily fetchable open access data, specifically two datasets on OpenNeuro (Markiewicz et al., 2021) with diverse participant profiles: *ds000228* (Richardson et al., 2019) and *ds000030* (Bilder et al., 2020). *ds000228* contains adult and child samples, and *ds000030* includes psychiatric conditions. The benchmark systematically evaluates the impact of denoising choices using a series of metrics based on past research (Ciric et al., 2017; Parkes et al., 2018).
- Our third objective was to turn this benchmark into a fully reproducible and interactive research object. We combined a series of technologies, including software containers (Gorgolewski et al., 2017), the Jupyter Book project (Granger & Perez, 2021), and the NeuroLibre preprint service (Karakuzu et al., 2022) in order to create the first fully reproducible benchmark of denoising strategies for fMRI resting-state connectivity.
- Our fourth and last objective was to demonstrate that our approach can be used to evaluate the robustness of the benchmark, by identifying possible differences across multiple versions of fMRIPrep.

## Results

### Software implementation

We designed two APIs for users to perform denoising of fMRI time series using Nilearn, based on fMRIPrep outputs. The APIs are maintainable, i.e., composed of modular and well-tested code, and user-friendly, i.e., the syntax is standard and robust to errors. The confounds are loaded by the APIs in a format compatible with downstream Nilearn analysis functions. The first, basic API retrieves different classes of confound regressors sorted in categories of noise, `nilearn.interfaces.fmriprep.load_confounds` (simplified as `load_confounds` in the following sections). The second, higher level API implements common strategies from the denoising literature, `nilearn.interfaces.fmriprep.load_confounds_strategy` (simplified as `load_confounds_strategy` in the following sections). The `load_confounds` and `load_confounds_strategy` APIs are available from Nilearn version 0.9.0 onwards. The following section describes both APIs in greater detail.

#### load_confounds: basic noise components

The following Python code snippet demonstrates the basic usage of load_confounds.

~~~
from nilearn.interfaces.fmriprep import load_confounds
confounds_simple, sample_mask = load_confounds(
      fmri_filenames,
      strategy=["high_pass", "motion", "wm_csf"],
      motion="basic", wm_csf="basic")
~~~

- **`fmri_filenames`:** path to processed image files, optionally as a list of paths.
- **`strategy`**: A list defining the categories of confound variables to use. Amongst the three in this example, `motion` and `wm_csf` are further tunable.
- **`motion`** and **`wm_csf`**: additional parameters with four options
  - `basic`: original parameters
  - `power2`: original parameters + quadratic terms
  - `derivatives`: original parameters + 1st temporal derivatives
  - `full`: original parameters + 1st temporal derivatives + quadratic terms + power2d derivatives

The `load_confounds` API fetches specific categories of confound variables, such as motion parameters. It is possible to fine-tune these categories through various options, such as the order of expansion of motion parameters. The implementation only supports fMRIPrep version 1.4 and above, and requires the fMRIPrep output directory in its original format. Users specify the path of a preprocessed functional file (file ending with `desc-preproc_bold.nii.gz` or `desc-smoothAROMAnonaggr_bold.nii.gz` in the case of ICA-AROMA). Warnings and errors inform the user if files or confounds were missing, for example if fMRIPrep was run without the option for ICA-AROMA yet users request ICA-AROMA confounds, or try to load an preprocessed fMRI output not suited for combination with ICA-AROMA regressors. The function returns the confound variables in a Pandas `DataFrame` object (McKinney, 2010; The pandas development team, 2023) and a time sample mask. The sample mask indexes the time points to be kept. The function can also be used with a list of input files, in which case it returns a list of confounds `DataFrames` and a list of time sample masks. A parameter called `strategy` can be used to pass a list of different categories of noise regressors to include in the confounds: `motion`, `wm_csf`, `global_signal`, `scrub`, `compcor`, `ica_aroma`, `high_pass`, `non_steady_state`. For each noise category, additional function parameters are available to tune the corresponding noise variables (please refer to Nilearn documentation^1^ for more details). See Annex A for a literature review and discussion for each category of common noise sources.

#### load_confounds_strategy: pre-defined strategies

The following code snippet demonstrates the basic usage of `load_confounds_strategy`. This snippet retrieves the same confounds variables as described in the example for `load_confounds`.

~~~
from nilearn.interfaces.fmriprep import load_confounds_strategy
confounds_simple, sample_mask = load_confounds_strategy(
      fmri_filenames,
      denoise_strategy="simple")
~~~

- **`fmri_filenames`:** path to processed image files, optionally as a list of paths.
- **`denoise_strategy`**: The name of a predefined strategy (see Table 1).

`load_confounds_strategy` provides an interface to select a complete set of curated confounds reproducing a common strategy used in the literature, with limited parameters for user customisation. There are four possible strategies that can be implemented from fMRIPrep confounds:

- `simple` (Fox et al., 2005): motion parameters and tissue signal
- `scrubbing` (Power et al., 2012): volume censoring, motion parameters, and tissue signal
- `compcor` (Behzadi et al., 2007): anatomical compcor and motion parameters
- `ica_aroma` (Pruim, Mennes, van Rooij, et al., 2015): ICA-AROMA based denoising and tissue signal

All strategies, except `compcor`, provide an option to add global signal to the confound regressors. The predefined strategies and associated noise components are listed in Table 1. Parameters that can be customized are indicated with a *. See the Nilearn documentation^2^ for more details. See Annex B for a more in-depth review of common denoising strategies in the literature and Annex C for a summary of evaluation benchmarks using these strategies.

**Table 1.** Correspondence of load_confounds parameters to predefined denoising strategies in load_confounds_strategy.

| Strategy | simple | scrubbing | compcor | ica_aroma |
| --- | --- | --- | --- | --- |
| high_pass | True | True | True | True |
| motion | full* | full* | full* | N/A |
| wm_csf | basic* | full | N/A | basic* |
| global_signal | None* | None* | N/A | None* |
| scrub | N/A | 5* | N/A | N/A |
| fd_threshold | N/A | 0.2* | N/A | N/A |
| std_dvars_threshold | N/A | 3* | N/A | N/A |
| compcor | N/A | N/A | anat_combined* | N/A |
| n_compcor | N/A | N/A | all* | N/A |
| ica_aroma | N/A | N/A | N/A | full |
| demean | True* | True* | True* | True* |

\* Parameters with customisable parameters.

### Denoising workflow

The denoising workflow is implemented through Nilearn. Figure 1 presents the graphic summary of the workflow. An fMRI dataset in the Brain Imaging Data Structure (BIDS) standard was first passed to fMRIPrep. Brain parcellation atlases were retrieved through the TemplateFlow (Ciric et al., 2022) Python client (see https://www.templateflow.org/usage/client/). In cases where an atlas was absent from TemplateFlow, it was converted into TemplateFlow naming convention to enable use of the Python client. Each atlas was passed to the `NiftiLabelsMasker` or `NiftiMapsMasker` for time series extraction. fMRIPrep outputs were input to a Nilearn-based connectome generating workflow using `load_confounds_strategy`. The filtered confounds and the corresponding preprocessed NIFTI images were then passed to the Nilearn masker generated with the atlas. The time series and connectomes were saved as the main outputs for further analysis.

**Figure 1.**
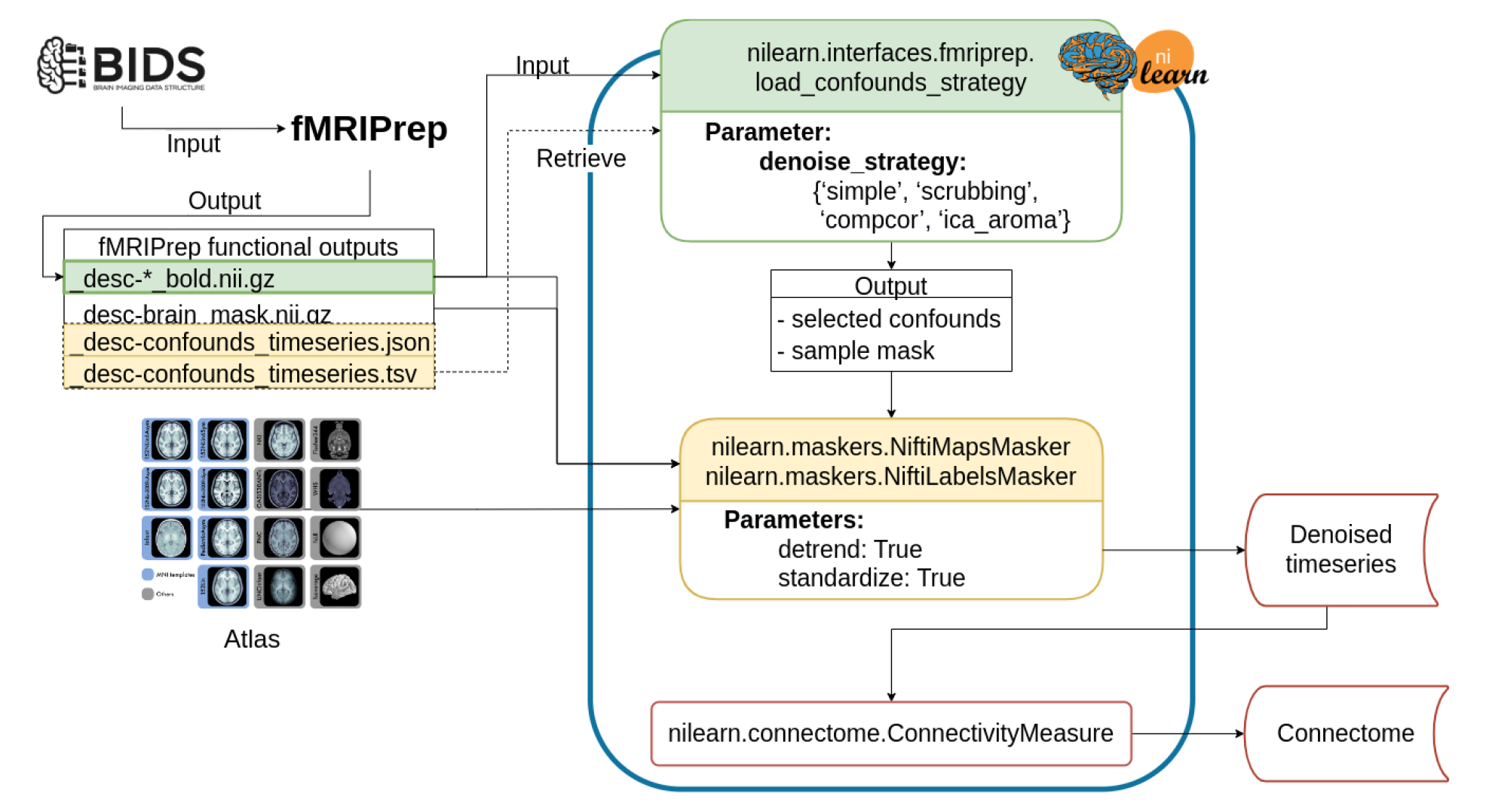
Workflow for post-fMRIPrep time series extraction with Nilearn tools. *The Python-based workflow describes the basic procedure to generate functional connectomes from fMRIPrep outputs with a Nilearn data loading routine (e.g., `NiftiMapsMasker` or `NiftiLabelsMasker)`, fMRIPrep confounds output retrieval function (e.g., `load_confounds_strategy)`, and connectome generation routine (`ConnectivityMeasure)`. Path to the preprocessed image data is passed to `load_confounds_strategy` and the function fetches the associated confounds from the .tsv file. The path of an atlas and the path of the preprocessed image file is then passed to the masker, along with the confounds, for time series extraction. The time series are then passed to `ConnectivityMeasure` for generating connectomes*.

### Benchmark workflow

OpenNeuro datasets were retrieved through DataLad (Halchenko et al., 2021) and fMRIPrep images were pulled from DockerHub. SLURM job submission scripts to process the fMRI data were generated with the Python tool fMRIPrep-SLURM (https://github.com/SIMEXP/fmriprep-slurm). The fMRIPrep derivatives and atlas retrieved from the TemplateFlow archive were passed to the connectome workflow described in Figure 1. We extracted the signals using a range of atlases at various resolutions (see Materials and Methods for details). For each parcellation scheme and each fMRI dataset, 10 sets of time series were generated, including one baseline and 9 different denoising strategies (see Table 2). We report the quality metrics and break down the effect on each dataset, preprocessed with fMRIPrep 20.2.1 long-term support branch (LTS). Motion characteristics were also generated per dataset and used to exclude fMRI runs with excessive motion from entering the benchmark. Trends in each atlas were similar, so we combined all atlases for the following report. The detailed breakdown by parcellation scheme can be found in the associated Jupyter Book (Wang et al., 2023). Figure 2 presents a graphical summary of the benchmark workflow.

**Figure 2.**
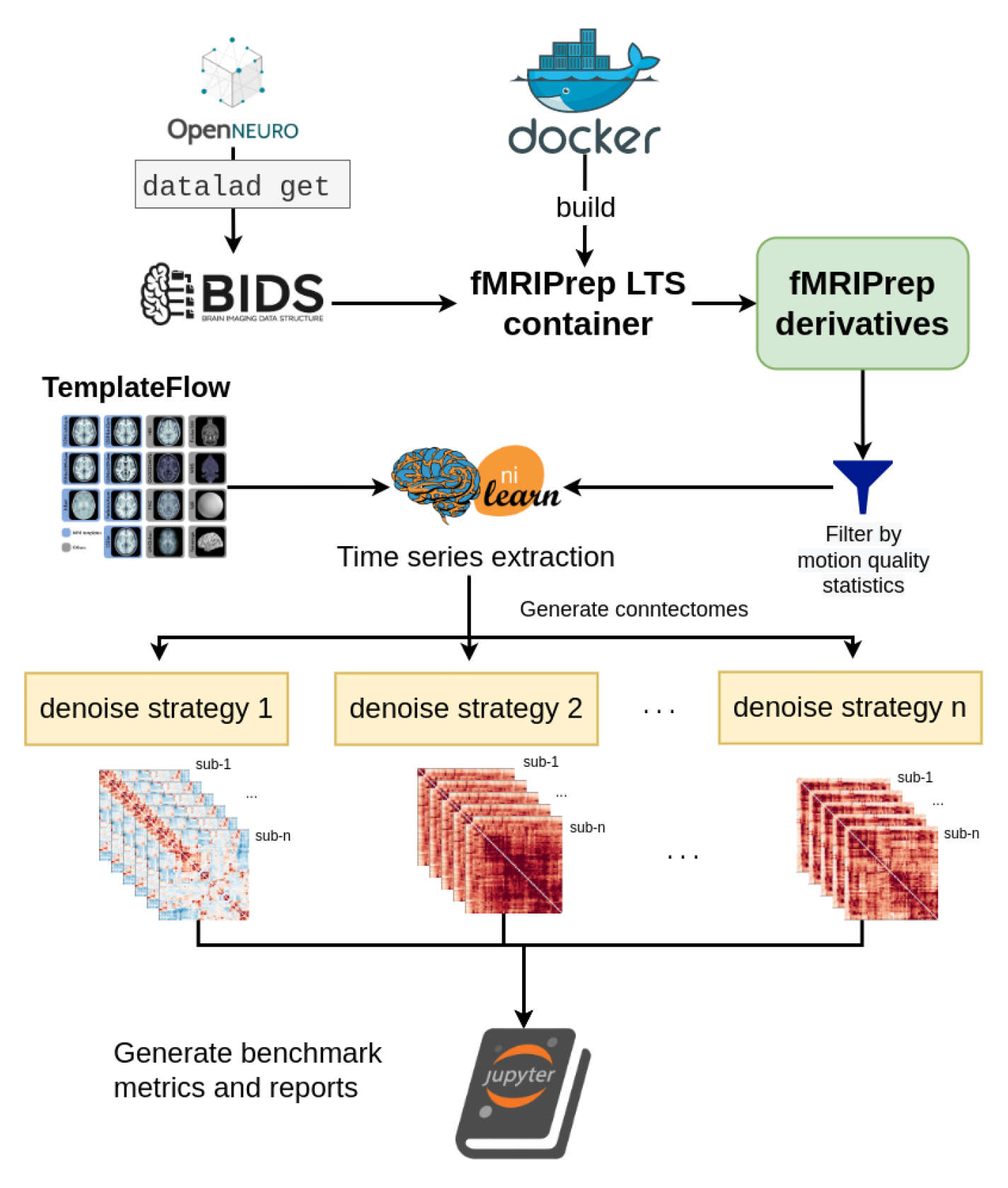
Denoising benchmark workflow. *The denoising benchmark workflow expands on the workflow in* *Figure 1* *(represented by the Nilearn logo in this figure). We retrieved the datasets from OpenNeuro through DataLad and all steps indicated with the arrows are implemented with bash scripts written for the SLURM scheduler. Atlases were either retrieved from the TemplateFlow archive or reformatted to fit the TemplateFlow format. The extracted time series, denoising metrics, and all metadata for generating the report are available on Zenodo (`10.5281/zenodo.6941757`)*.

**Table 2.** Strategies examined in the benchmark and associated parameters applied to load_confounds.

| strategy | image | high_pass | motion | wm_csf | global_signal | scrub | fd_thresh (mm) | compcor mask | n_compcor | ica_aroma | demean |
| --- | --- | --- | --- | --- | --- | --- | --- | --- | --- | --- | --- |
| baseline | desc-prep<br>roc_bold | True | N/A | N/A | N/A | N/A | N/A | N/A | N/A | N/A | True |
| simple | desc-prep<br>roc_bold | True | full | basic | N/A | N/A | N/A | N/A | N/A | N/A | True |
| simple+gsr | desc-prep<br>roc_bold | True | full | basic | basic | N/A | N/A | N/A | N/A | N/A | True |
| scrubbing.5 | desc-prep<br>roc_bold | True | full | full | N/A | 5 | 0.5 | N/A | N/A | N/A | True |
| scrubbing.5+gsr | desc-prep<br>roc_bold | True | full | full | basic | 5 | 0.5 | N/A | N/A | N/A | True |
| scrubbing.2 | desc-prep<br>roc_bold | True | full | full | N/A | 5 | 0.2 | N/A | N/A | N/A | True |
| scrubbing.2+gsr | desc-prep<br>roc_bold | True | full | full | basic | 5 | 0.2 | N/A | N/A | N/A | True |
| compcor | desc-prep<br>roc_bold | True | full | N/A | N/A | N/A | N/A | anat_combined | all* | N/A | True |
| compcor6 | desc-prep<br>roc_bold | True | full | N/A | N/A | N/A | N/A | anat_combined | 6 | N/A | True |
| aroma** | desc-smoothAROMAno<br>naggr_bold | True | N/A | basic | N/A | N/A | N/A | N/A | N/A | full*** | True |

\* 50% variance explained.
\*\* In Ciric et al. (2017), there was a variation of the ICA-AROMA strategy including global signal regressor. The global signal regressor generated by fMRIPrep does not follow the recommendation of Pruim et al. (2015). The result of ICA-AROMA+GSR can be found in the first version of the preprint.
\*\*\* Referring to the non-aggressive implementation in Pruim and colleagues' work (2015)

### Benchmark results from fMRIPrep 20.2.1 LTS

We reported the demographic information and the gross mean framewise displacement before and after excluding subjects with high motion. We then aimed to assess the overall similarity between connectomes generated from each denoising strategy, and evaluated the denoising strategies using four metrics from Ciric and colleagues’ benchmark (2017):

1. Loss of degrees of freedom: sum of number of regressors used and number of volumes censored.
2. Quality control / functional connectivity (QC-FC; Power et al., 2015): partial correlation between motion and connectivity with age and sex as covariates.
3. Distance-dependent effects of motion on connectivity (DM-FC Power et al., 2012): correlation between node-wise Euclidean distance and QC-FC.
4. Network modularity (Satterthwaite et al., 2012): graph community detection based on Louvain method, implemented in the Brain Connectome Toolbox.

#### Significant differences in motion levels existed both between datasets, and within-dataset, across clinical and demographic subgroups

We applied a motion threshold to exclude subjects with marked motion in the two OpenNeuro datasets: dataset *ds000228* (N=155) (Richardson et al., 2019) and dataset *ds000030* (N=212) (Bilder et al., 2020). Table 3 shows the demographic information of subjects in each dataset after the automatic motion quality control. Following this, we checked the difference in the mean framewise displacement of each sample and the sub-groups (Figure 3). In *ds000228*, there was still a significant difference (t(73) = −2.17, p = 0.033) in motion during the scan captured by mean framewise displacement between the child (M = 0.17, SD = 0.05, n = 51) and adult samples (M = 0.15, SD = 0.04, n = 24). In *ds000030*, the only patient group that showed a difference compared to control subjects (M = 0.12, SD = 0.04, n = 88) was the schizophrenia group (M = 0.16, SD = 0.05, n = 19; t(105) = −3.49, p = 0.033). There was no difference between the control and ADHD group (M = 0.12, SD = 0.05, n = 32; t(118) = 0.04, p = 0.966), or the bipolar group (M = 0.13, SD = 0.05, n = 29; t(115) = −1.24, p = 0.216). In summary, children moved more than adults, and subjects with schizophrenia moved more than controls.

**Figure. 3.**
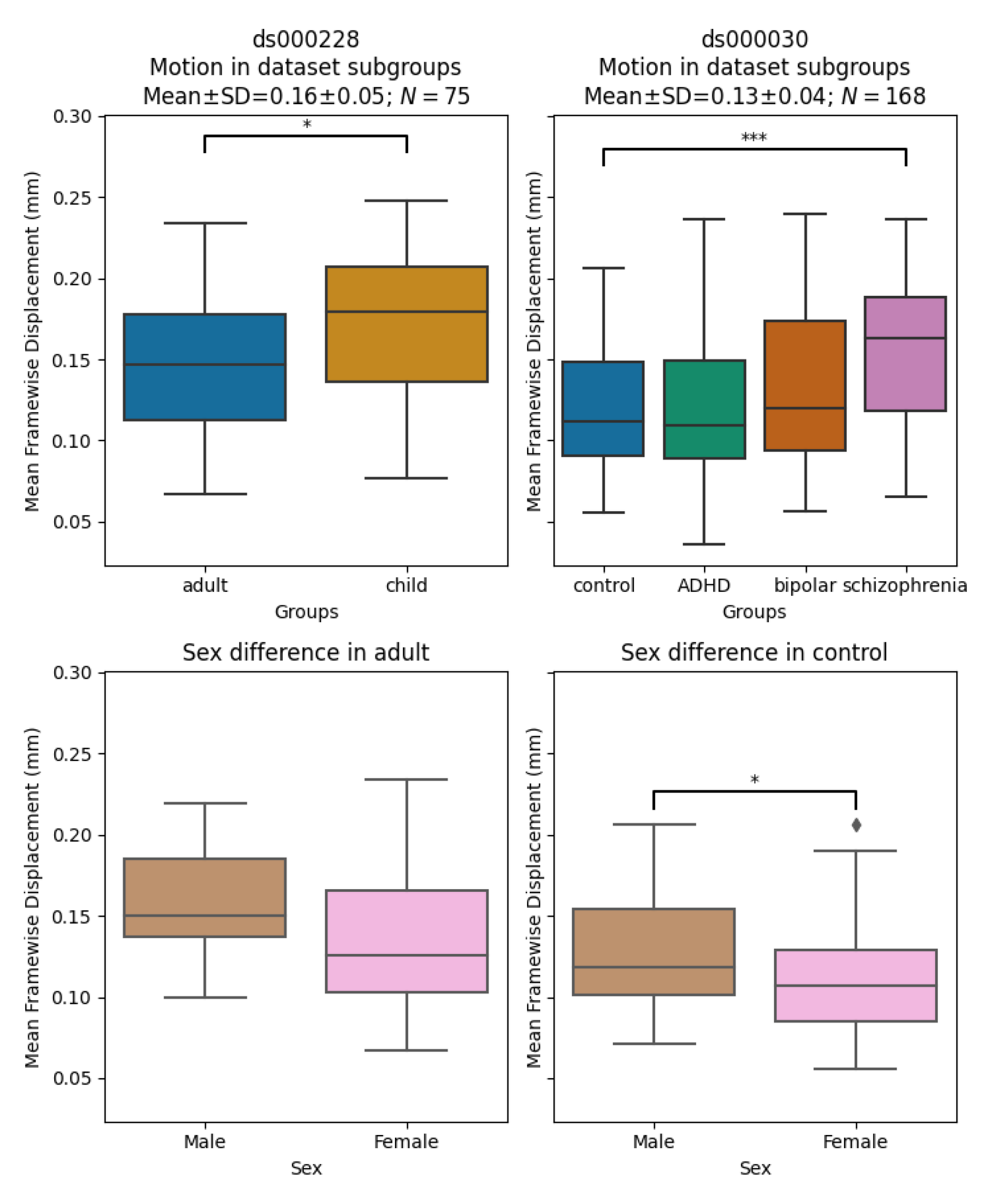
Mean framewise displacement of each dataset. *To evaluate the metrics in a practical analytic scenario, we excluded subjects with high motion while preserving 1 minute of data for functional connectivity calculation: gross mean framewise displacement > 0.25 mm, above 80.0% of volumes removed while scrubbing with a 0.2 mm threshold. In ds000228, the child group still had higher motion compared to the adult groups. In ds000030, where all subjects were adults, the control group only showed significant differences in motion with the schizophrenia group. In both datasets, the sample sizes from each group were highly imbalanced (see Table 3), hence no between group differences were assessed in quality metrics analysis*.

**Table 3.**
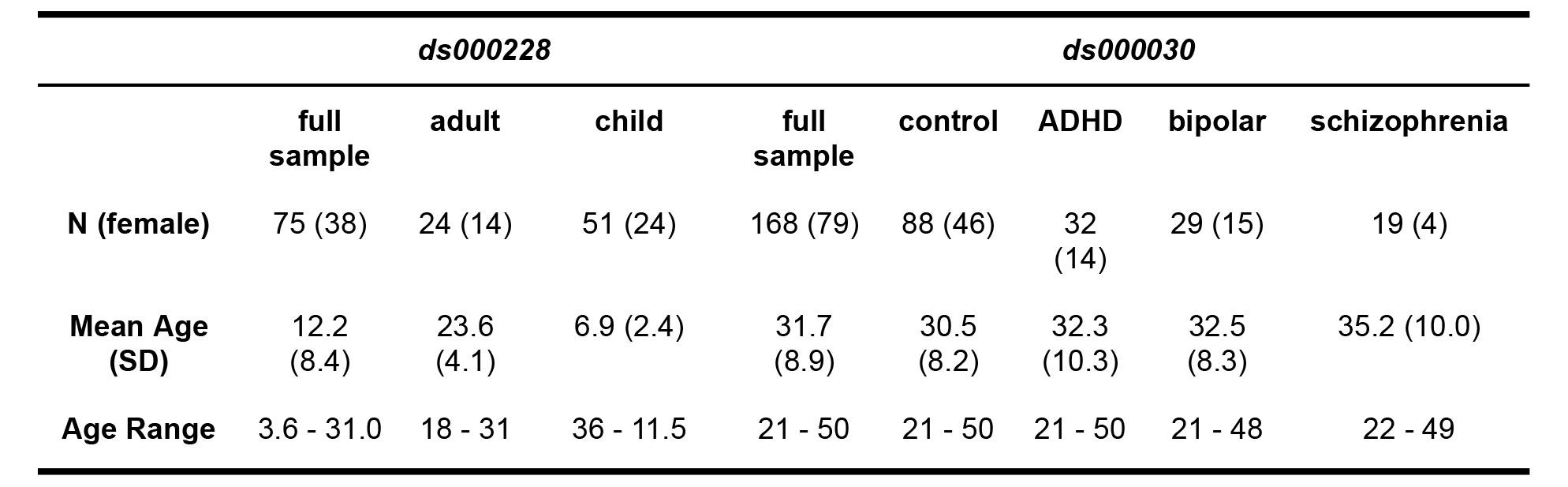
Sample demographic information after removing subjects with high motion.

|  | *ds000228* |  |  | *ds000030* |  |  |  |  |
| --- | --- | --- | --- | --- | --- | --- | --- | --- |
|  | full sample | adult | child | full sample | control | ADHD | bipolar | schizophrenia |
| **N (female)** | 75 (38) | 24 (14) | 51 (24) | 168 (79) | 88 (46) | 32 (14) | 29 (15) | 19 (4) |
| **Mean Age (SD)** | 12.2 (8.4) | 23.6 (4.1) | 6.9 (2.4) | 31.7 (8.9) | 30.5 (8.2) | 32.3 (10.3) | 32.5 (8.3) | 35.2 (10.0) |
| **Age Range** | 3.6 - 31.0 | 18 - 31 | 36 - 11.5 | 21 - 50 | 21 - 50 | 21 - 50 | 21 - 48 | 22 - 49 |

<sup>3</sup> https://preprint.neurolibre.org/10.55458/neurolibre.00012/

We also examined the differences between male and female in the control groups of the two datasets: the adult sample for *ds000228* and healthy control for *ds000030*. In *ds000228*, we found no significant differences (male: M = 0.16, SD = 0.04; female: M = 0.14, SD = 0.05; t(22) = 1.19, p = 0.249). In *ds000030* we found the male sample (M = 0.13, SD = 0.04) showed higher mean framewise displacement than the female sample (M = 0.11, SD = 0.04; t(86) = 2.17, p = 0.033).

Due to the imbalanced samples per group and low number of subjects in certain groups after the automatic motion quality control, we collapsed all groups within each dataset to avoid speculation on underpowered samples in the results. For a breakdown of each metric by atlas, please see the supplemental Jupyter Book^3^.

### Most denoising strategies converged on a consistent average connectome structure

With the benchmark workflow in place, we first aimed to assess the overall similarity between connectomes generated from each denoising strategy. We calculated Pearson’s correlations between connectomes generated from all strategies presented in the benchmark (Figure 4). The connectome correlation pattern across denoising strategies was similar in both datasets. Overall, the strategies displayed at least moderate similarity with each other, with Pearson’s correlations above 0.6. There were two large clusters of highly-related strategies, driven by the presence (or lack) of global signal regression. Within each cluster of strategies, the correlations amongst the strategies were strong, with values above 0.9. `baseline` and `aroma` did not fit well in either of the two clusters, indicating that denoising generally impacts the connectome structure, and that the ICA-AROMA might be sensitive to different sources of noise, compared to those captured by other strategies in the benchmark.

**Figure 4.**
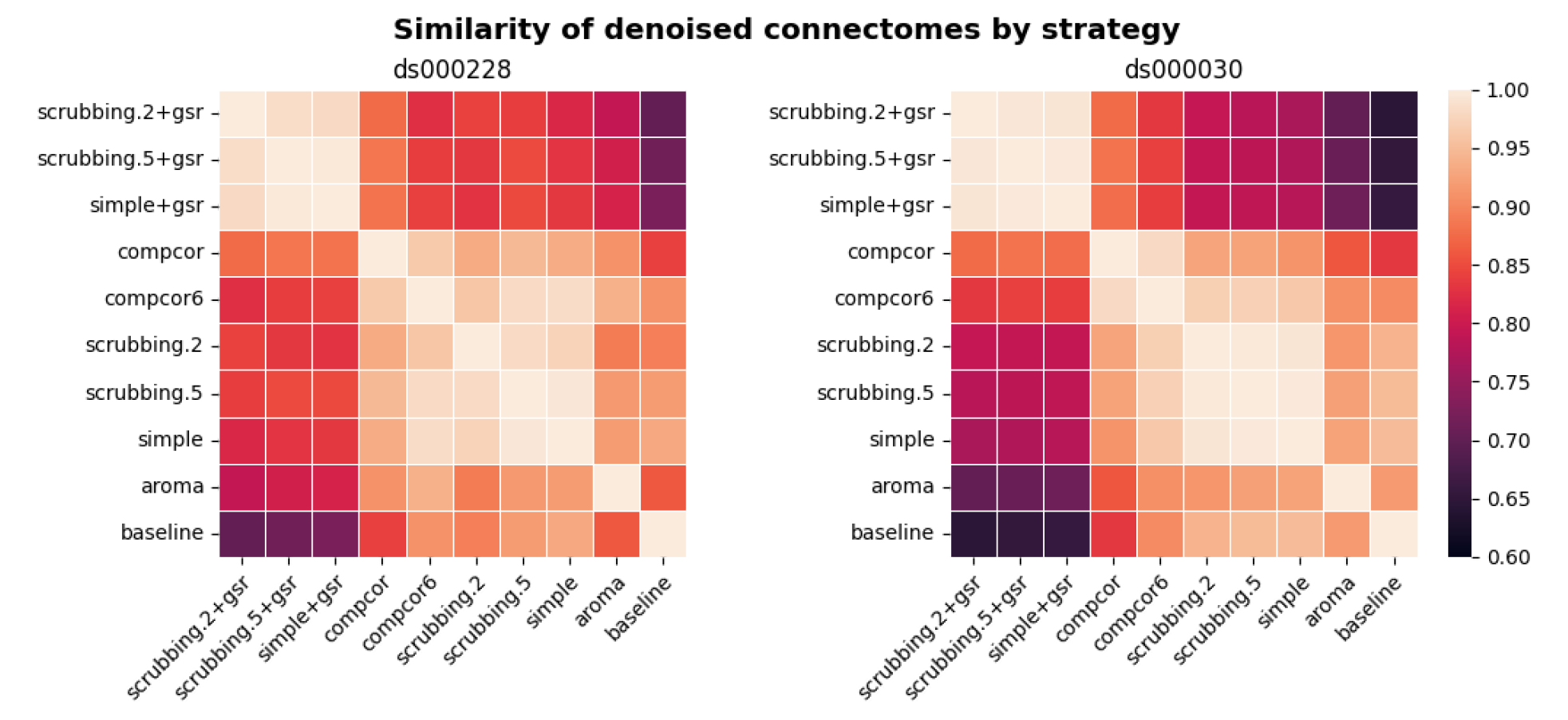
Similarity of denoised connectomes. *For each parcellation scheme, we computed a correlation matrix across connectomes generated with the ten strategies. These correlation matrices were then averaged across the parcellation schemes within each dataset. Two large clusters of strategies emerged: with versus without global signal regression, with fairly high similarity in connectomes within each cluster*.

#### Loss in temporal degrees of freedom varied markedly across strategies and datasets

In previous research, the loss of temporal degrees of freedom has shown an impact on the subsequent data analysis. Higher loss in temporal degrees of freedom can spuriously increase functional connectivity (Yan et al., 2013). Volume censoring-based and data-driven strategies (ICA-AROMA and some variations of CompCor) introduce variability to degrees of freedom and can bias group level comparisons (Ciric et al., 2017).

The loss of temporal degrees of freedom is the sum of the number of regressors used and censored volume lost. Depending on the length of the scan, the number of discrete cosine-basis regressors can differ given the same repetition time (TR). The two datasets we analyzed contain different numbers of discrete cosine-basis regressors (*ds000228*: 4; *ds000030*: 3) due to difference in time series length (*ds000228*: 168; *ds000030*: 152). The `simple` and `simple+gsr` strategies include the same amount of head motion and tissue signal regressors between the two datasets (`simple`: 26, `simple+gsr`: 27). For volume censoring strategies, we observed a higher loss in volumes in *ds000228*, compared to *ds000030*. (number of excised volumes at 0.5 mm: *ds000030*: 2.5(4.4) range=[0 21], *ds000228*: 9.3(8.8) range=[0 30]; number of excised volumes at 0.2 mm: *ds000030*: 29.4(30.1) range=[0 110], *ds000228*: 53.0(34.1) range=[1 130]. `compcor` also showed variability in numbers of regressors when using all components that explain 50% of signal variance (number of CompCor regressors: *ds000030*: 47.9(3.9) range=[35 54], *ds000228*: 42.5(8.9) range=[5 58]). ICA-AROMA regressors in strategy `aroma` showed variability in numbers of regressors (number of ICA-AROMA regressors: *ds000030*: 16.0(4.6) range=[6 29], *ds000228*: 20.9(6.3); range=[7 38]). The average loss in temporal degrees of freedom is summarized in Figure 5.

**Figure 5.**
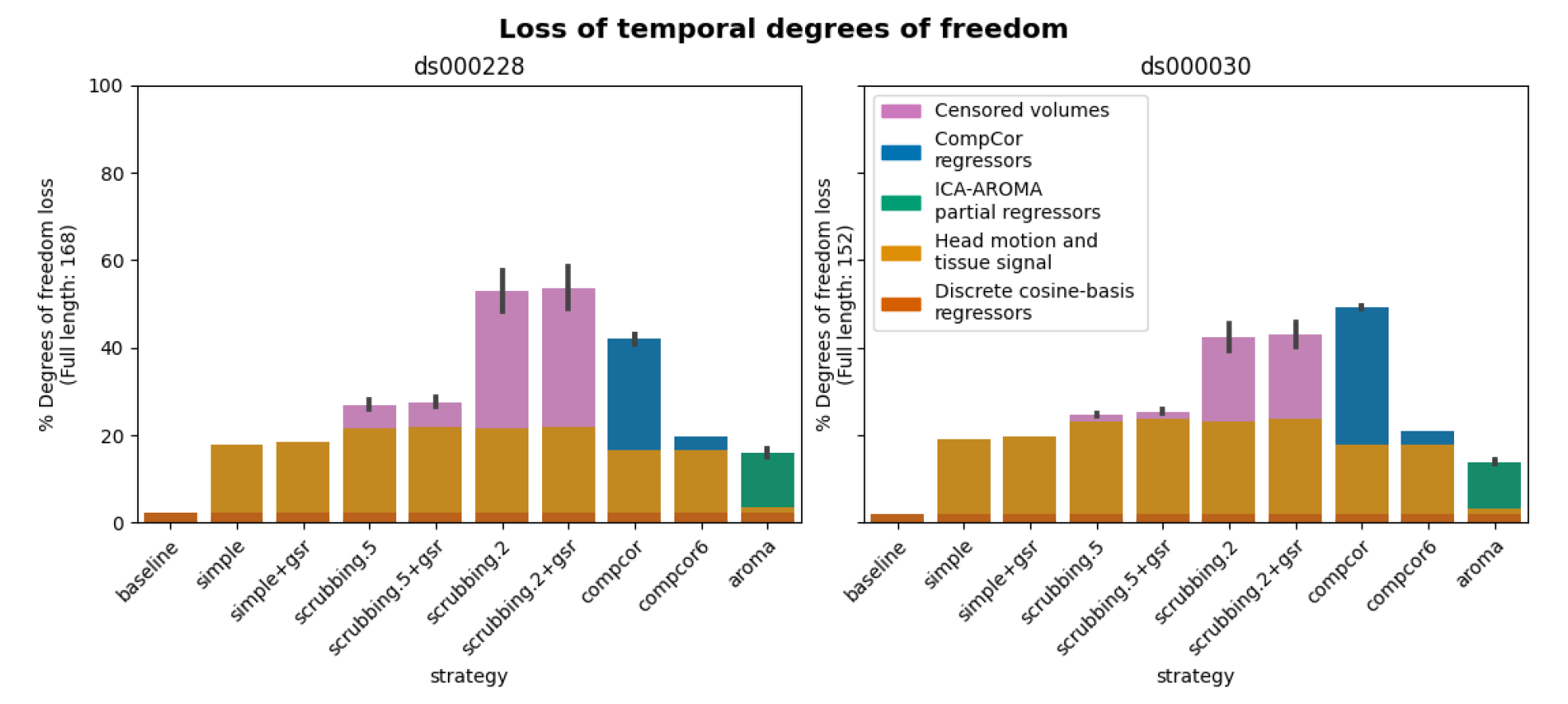
Percentage of loss in temporal degrees of freedom according to strategy and dataset. Bars show the average percentage of the number of regressors to the length of the scan amongst all subjects. Error bars indicate 95% confidence interval. The two datasets contain different numbers of discrete cosine-basis regressors (*ds000228*: 4; *ds000030*: 3). *`Compcor` (anatomical CompCor extracted from a WM/CSF combined map, cut off at 50% variance) and ICA-AROMA-based strategies (`aroma`) show variability depending on the number of noise components detected*.

The loss of degrees of freedom per strategy varied across the two datasets shown in the benchmark. The two datasets showed different loss of degrees of freedom in scrubbing-based strategies, while using the same gross motion-based exclusion criterias. This was expected, as the amount of motion between time points was higher in ds000228, a dataset consisting mostly of children. `compcor` did not always have a lower loss of degrees of freedom in *ds000030*. `aroma` had the lowest loss of temporal degrees of freedom in *ds000030*.

### Quality control / functional connectivity (QC-FC) showed a heterogeneous impact of denoising strategies based on data profile

The denoising methods should aim to reduce the impact of motion on the data. To quantify the remaining impact of motion in connectomes, we adopted a metric proposed by Power and colleagues (2015) named quality control / functional connectivity (QC-FC). QC-FC is a partial correlation between mean framewise displacement and functional connectivity, with age and sex as covariates. Significance tests associated with the partial correlations were performed. P-values above the threshold of α = 0. 05 were deemed significant.

Scrubbing-based strategies consistently performed better than the baseline in both datasets. In *ds000228*, the most effective method according to QC-FC was `scrubbing.5` (scrubbing at a liberal threshold), followed by `scrubbing.2` and `simple`. All the GSR counterparts of the methods had slightly higher residual motion. Amongst all the data-driven methods, `compcor` performed the best. `compcor6` and `aroma` performed close to baseline. In *ds000030*, the best performing method was `compcor`, followed by `scrubbing.2` (aggressive scrubbing). The `simple` and `scrubbing.5` methods performed similarly as very few volumes were censored with a liberal threshold, and the GSR variations (`simple+gsr` and `scrubbing.5+gsr`) performed better than baseline (see Figure 6). `simple` performed close to the `baseline` in terms of the number of edges correlated with motion. The `aroma` and `compcor6` strategies were better than baseline. The average percentage of significant QC-FC and the average median of absolute value of QC-FC are presented in Figure 6 and Figure 7. In summary, based on a QC-FC evaluation, diverse strategies performed quite differently based on the dataset used for evaluation.

**Figure 6.**
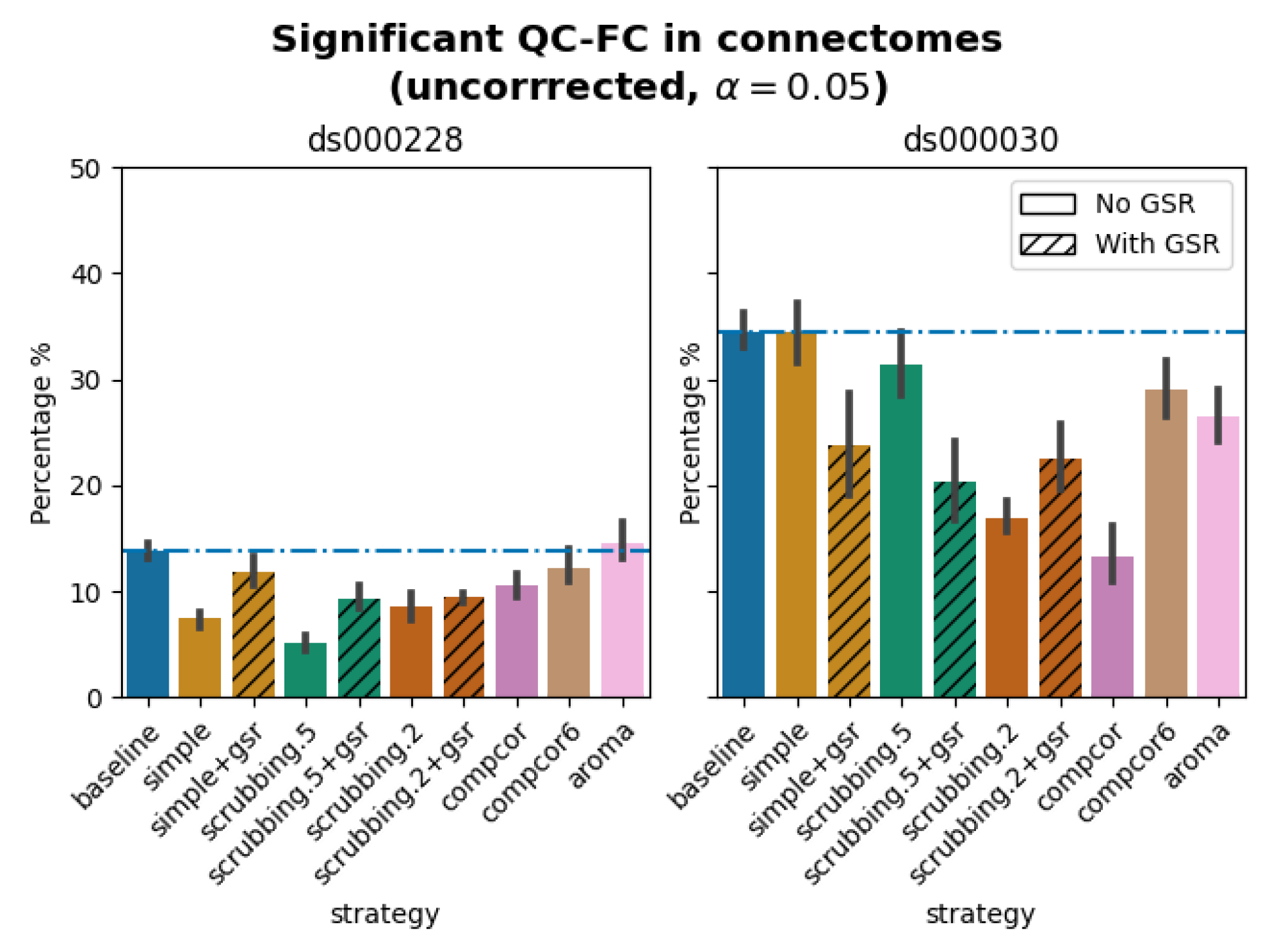
Significant QC-FC in connectomes. *Average percentage of edges significantly correlated with mean framewise displacement are summarized across all atlases as bar plots. Error bars represent the 95% confidence intervals of the average. The horizontal line represents the baseline. A lower percentage indicates less residual effect of motion after denoising on connectome edges. Significant QC-FC associations were detected with p<0.05, uncorrected for multiple comparisons. A version of the figure using false-discovery-rate correction for multiple comparisons can be found in supplemental Jupyter Book*.

**Figure 7.**
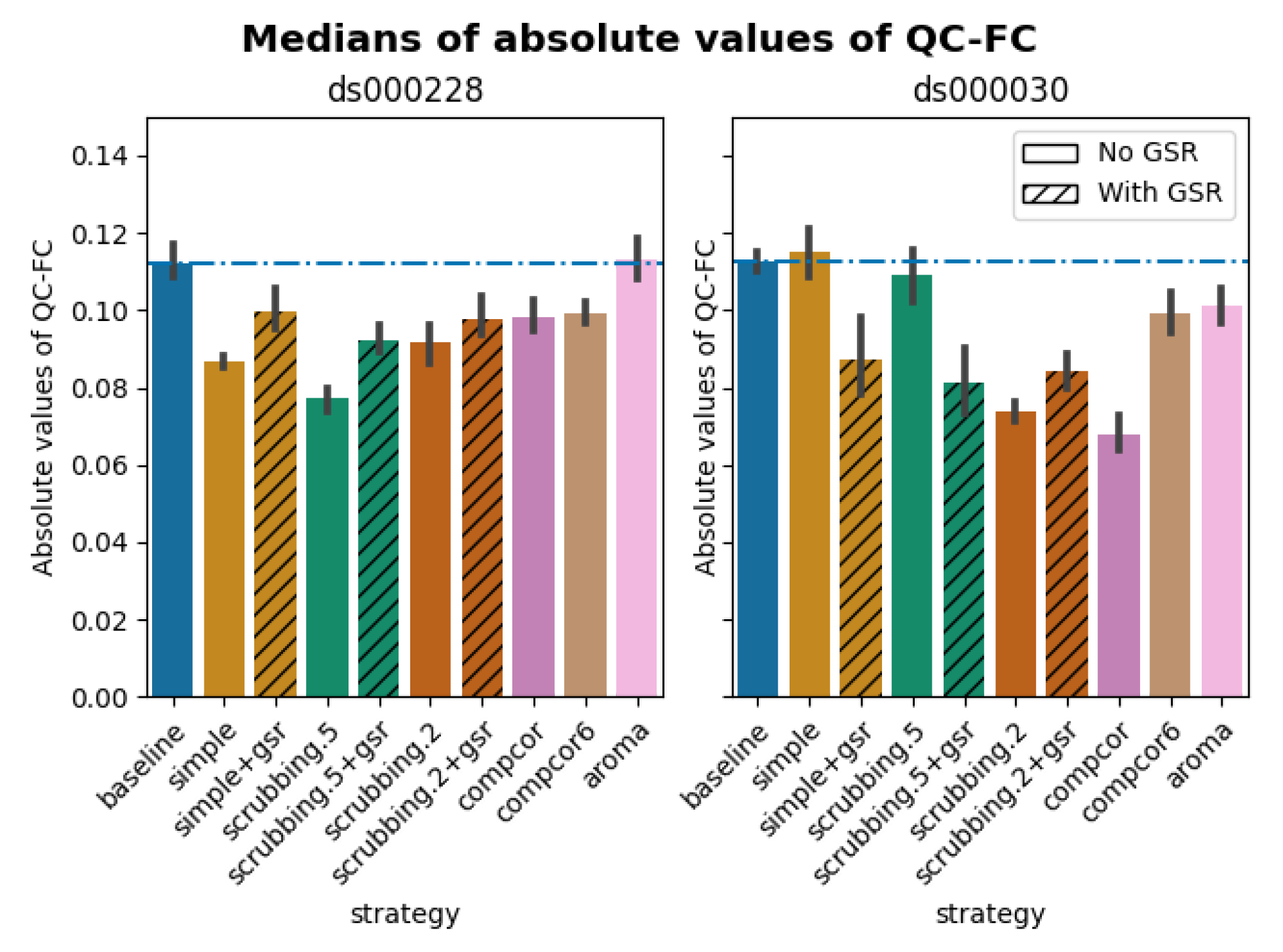
Medians of absolute values of QC-FC. *Median of absolute value of QC-FC, averaged across all atlases of choice. Error bars represent the confidence intervals of the average at 95%. Low absolute median values indicate less residual effect of motion after denoising. The horizontal line represents the baseline. Results observed with absolute QC-FC values are consistent with the percentage of edges with significant QC-FC associations, as reported in* *Figure 6*.

### Scrubbing-based strategies decreased distance-dependent effects of motion

The impact of motion on functional connectivity has been reported to be higher for brain parcels closer to each other in space (Power et al., 2012). To determine the residual distance-dependent effects of subject motion on functional connectivity (DM-FC), we calculated a correlation between the Euclidean distance between the centers of mass of each pair of parcels (Power et al., 2012) and the corresponding QC-FC correlations. We reported the absolute DM-FC correlation values and expected to see a general trend toward zero correlation after denoising.

All strategies performed better than the baseline in both datasets (Figure 8). We observed a trend consistent across both datasets, whereby strategies `scrubbing.2` and `scrubbing.2+gsr` were the most effective in reducing the correlation. `aroma` also performed consistently well in both datasets, ranked after `scrubbing.2`. In *ds000228*, `simple` was the least effective strategy for reducing distance dependency. Data-driven methods showed similar results to each other. `scrubbing.5` and `simple` greatly benefited from adding GSR in the regressors. In *ds000030*, the difference between `scrubbing.2` and other strategies was bigger than in *ds000228*, with the remainder performed similarly with each other. The impact of GSR was small with the exception of `scrubbing.2+gsr`. In summary, we observed similar trends across strategies between the two datasets, yet with differences in the magnitude of correlations. All strategies reduced the correlation lower than the baseline. Consistent with the literature, scrubbing strategies were the best at reducing distance dependency.

**Figure 8.**
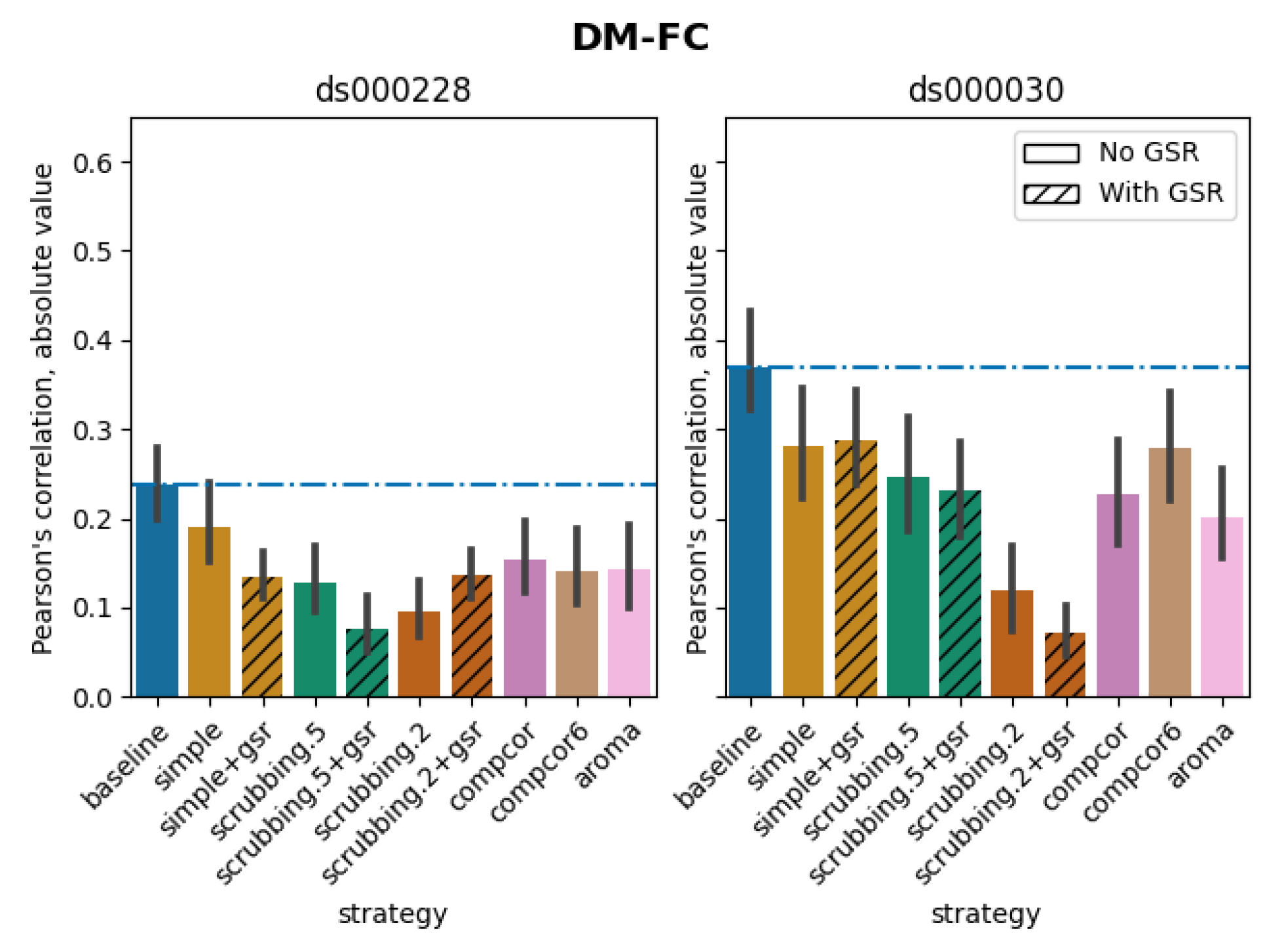

*Residual distance-dependent effects of subject motion on functional connectivity Average of absolute value of Pearson’s correlation between the Euclidean distance between node pairs and QC-FC, indicating distance-dependent of motion after denoising. A value closer to zero indicates less residual effect of motion after denoising. Error bars represent the standard deviation. The horizontal line represents the baseline. Strategies scrubbing.2 and scrubbing.2+gsr were the most effective in reducing the correlation in both datasets*.

### Global signal regression increases network modularity

Confound regressors have the harmful potential to remove real signals of interest as well as motion-related noise. To evaluate this possibility, we examined the impact of denoising strategies on a common graph feature, network modularity, generally regarded as a key feature of biological network organization (Satterthwaite et al., 2012). Network modularity was quantified using the Louvain method for graph community detection (Rubinov & Sporns, 2010). We computed the partial correlation between subjects’ modularity values and mean framewise displacement, using age and sex as covariates, following the implementation of Power and colleagues (2015).

The inclusion of global signal regressors increased average Louvain network modularity in both datasets (Figure 9, top panel). The remaining strategies performed as follows in both datasets, from best to worst: `compcor`, `scrubbing.2`, `scrubbing.5`, `simple`, `compcor6`, and `aroma`. In both datasets, `aroma` performed almost at the similar level as the `baseline`. We found fixed results in the ability of denoising in reducing the impact of motion on modularity (Figure 9 lower panels). In *ds000228*, we see `simple` and `scrubbing.5` reducing the impact of motion. In *ds000030*, only `scrubbing.2` performed better than baseline. In both datasets, the data-driven strategies and strategies with GSR performed consistently worse than `baseline`. The overall trend across strategies is similar to QC-FC with the exception of the `baseline` strategy (see Figure 6 and 7). The reason behind this observation could be a reduction of variance in the Louvain network modularity metric for GSR-based denoising strategies. We plotted the correlations of `baseline`, `scrubbing.2`, `scrubbing.2+gsr` from one parcellation scheme (DiFuMo 64 components) from *ds000030* to demonstrate this lack of variance (see Figure 10).

**Figure 9.**
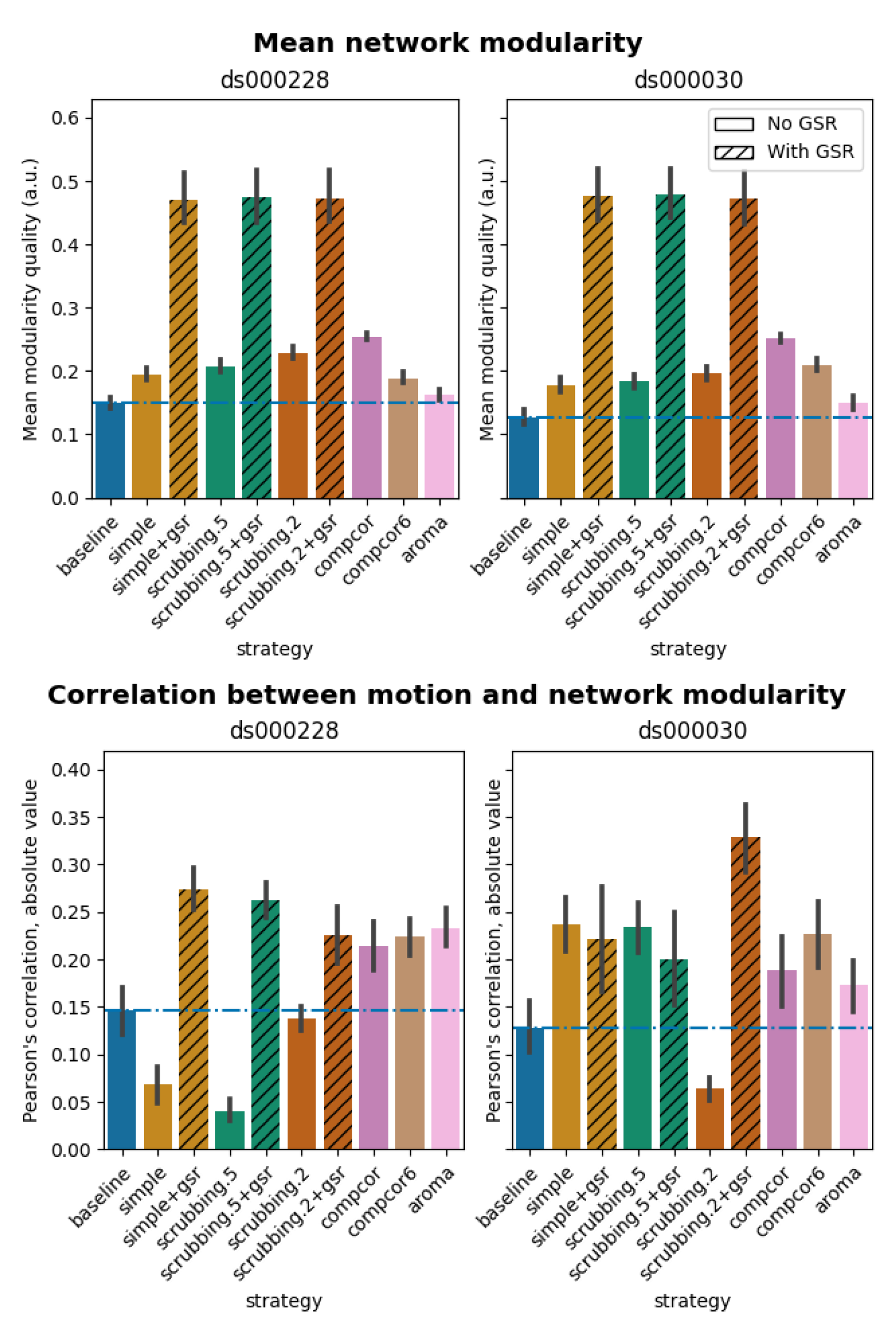
Network modularity measures. *Top: Average Louvain network modularity of all connectomes after denoising. Error bars represent the standard deviation. The horizontal line represents the baseline. In both datasets, strategies including the global signal regressor(s) have higher modularity values. Bottom: Average Pearson’s correlation between mean framewise displacement and Louvain network modularity after denoising. A value closer to zero indicates less residual effect of motion after denoising*.

**Figure 10.**
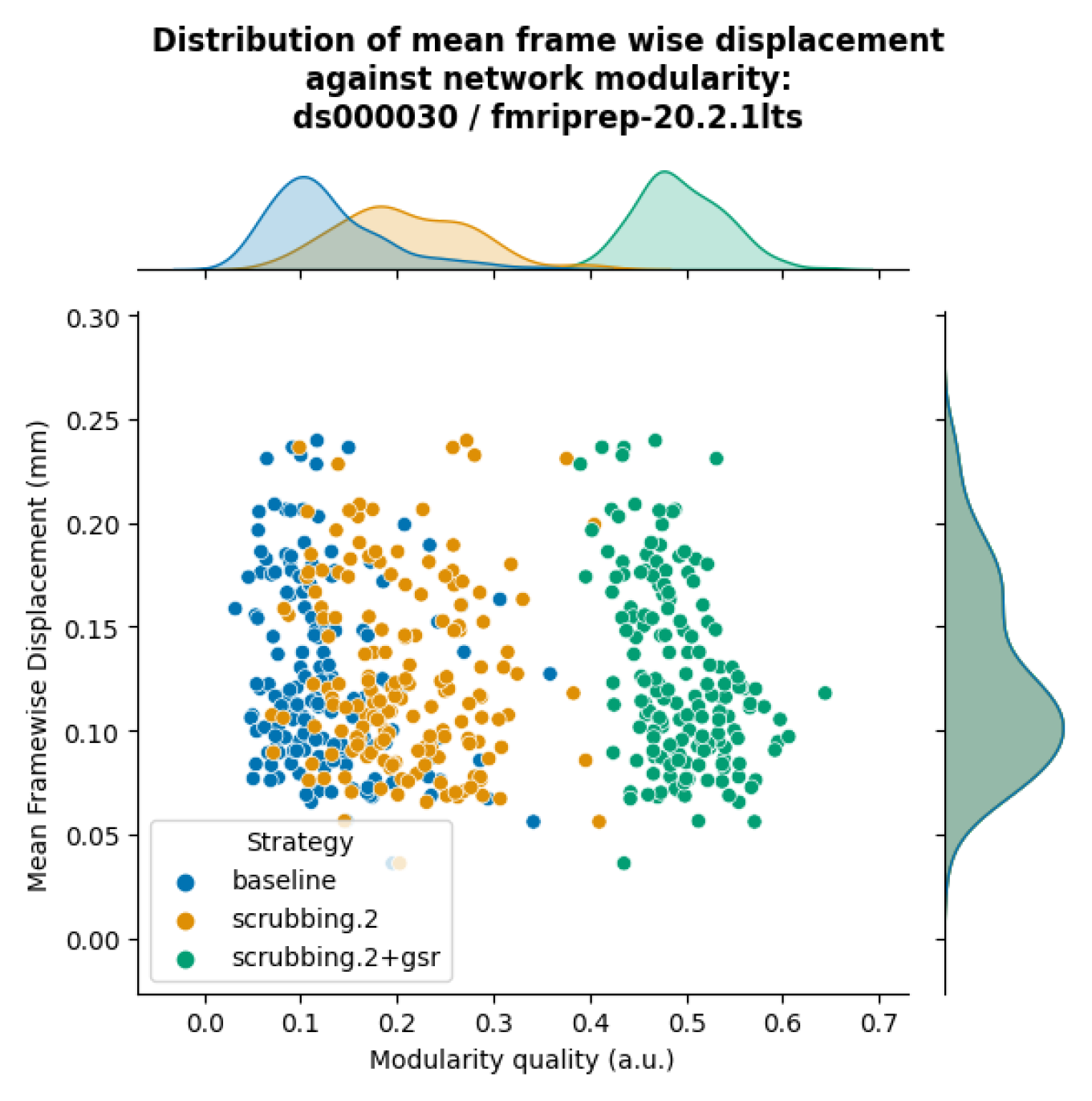
Correlation between mean framewise displacement and Louvain network modularity after denoising. *We observed a lack of variance in Louvain network modularity, and shrinkage of the distribution with the inclusion of GSR. Due to the lack of variability, assessing residual motion in network modularity might not be a good metric to evaluate the quality of connectivity data*.

### Data-driven denoising strategies showed inconsistent evaluation outcomes between two fMRIPrep versions

Different versions of the same software could produce differences in the outcomes of our denoising evaluation. To gain insight into the stability of fMRIPrep, we examined whether a few key observations from fMRIPrep 20.2.1 LTS remained salient in fMRIPrep 20.2.5 LTS, specifically:

1. High loss of temporal degrees of freedom for `scrubbing.2` in *ds000228* and compcor for *ds000030*.
2. `aroma` performed close to `baseline` in QC-FC for *ds000228*.
3. `simple` performed close to `baseline` in QC-FC for *ds000030*.
4. `scrubbing.2` and `scrubbing.2+gsr` were the best strategies to reduce DM-FC.
5. GSR-enabled strategies showed higher network modularity.

Observations 1, 4, and 5 from 20.2.5 LTS were consistent with results from 20.2.1 LTS. The results of QC-FC demonstrated similar overall trends in 20.2.5 LTS, but with `aroma` performing worse than `baseline` for *ds000228* (observation 2) and `simple` performing better than baseline for *ds000030* (observation 3) (see Figure 11). Inconsistency in outcomes across the two fMRIPrep versions were found in strategies with data-driven noise components. In version 20.2.5 LTS, and unlike 20.2.1 LTS, `comcpor6` performed worse than the `baseline` in metric QC-FC for both datasets. In *ds000228*, `aroma` was the second worst performing strategy. For *ds000030*, the strategies with no data-driven noise components showed better performance in 20.2.5 LTS (Figure 11) than 20.2.1 LTS (see Figure 6).

**Figure 11.**
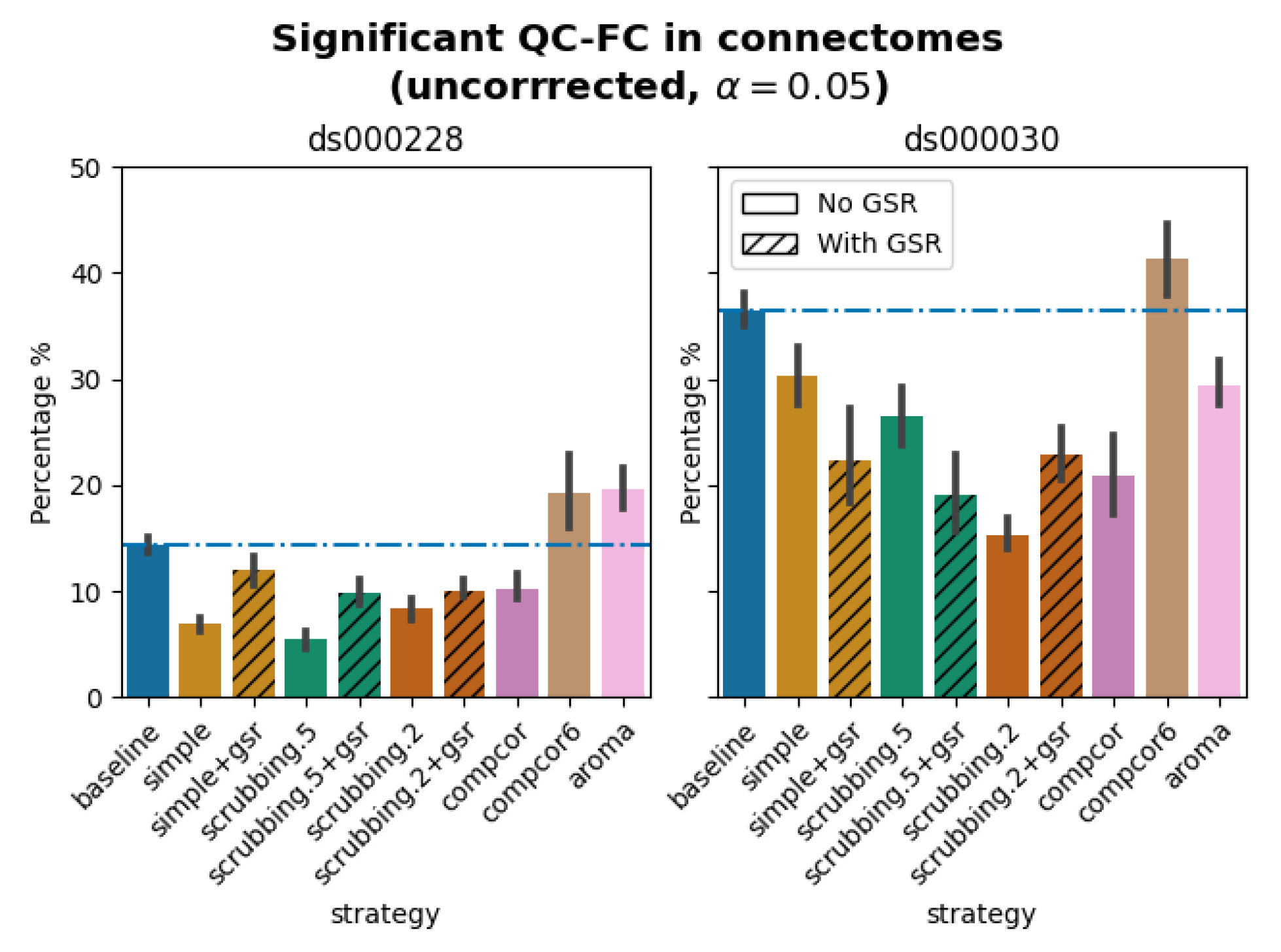
Significant QC-FC in connectomes compiled from 20.2.5 LTS. *Average percentage of edges significantly correlated with mean framewise displacement are summarized across all atlases as bar plots. Error bars represent the 95% confidence intervals of the average. The horizontal line represents the baseline. Lower values indicate less residual effect of motion after denoising. Data-driven denoising strategies showed inconsistent patterns compared to the same metric generated from 20.2.1 LTS outputs (Figure 6)*.

## Discussion

We aimed to create a re-executable benchmark to provide guidelines and accessible tools for denoising resting state functional connectivity data. The re-executable benchmark showed most denoising strategies, such as scrubbing-based strategies, `simple`, and strategies with GSR, performed in line with the literature. `aroma` showed an advantage in low degrees of freedom lost, while only stood out in DM-FC amongst all quality metrics. The metric performed consistently across the software versions with a marked exception in the data-driven denoising strategies (`compcor6`, `aroma`). This result demonstrates the necessity of distributing an executable research object for methods development and software testing, and providing accurate guidelines to users over time.

### The load_confounds and load_confounds_strategy APIs

The standardized APIs `load_confounds` and `load_confounds_strategy` are the core elements of the re-executable denoising benchmark. The APIs provide an easy way to implement classic denoising strategies from the literature, and can reduce the effort required, as well as errors, when using these strategies. Having clear and concise code also facilitates re-use and sharing of the denoising strategy used in a particular study, which improves reproducibility of science.

The new APIs developed for this project have been integrated in an established, popular software library, Nilearn(Abraham et al., 2014). The implementation of these APIs required other contributions to Nilearn and introduced new modules, in order to streamline the compatibility between the APIs and other data processing utilities. Specifically, we introduced a new module `nilearn.interfaces` dedicated to interacting with other neuroimaging software libraries and BIDS. We refactored the parameter `sample_mask` in all `masker` modules to allow volume censoring in the `signal.clean` function^4^. The `masker` modules implement a series of methods to convert 3D or 4D neuroimaging data into numerical arrays, for example extracting average time series from a brain parcellation. As a result, the outputs from `load_confounds` and `load_confounds_strategy`, as well as volume censoring information, can be directly ingested into all Nilearn masker objects. Thanks to these contributions, it is now possible to construct a complete Python-based fMRIPrep post-processing workflow with very concise code. Documentation for this workflow can be found in the Nilearn User Guide library^5^, and users can adapt code from the Nilearn tutorial to implement denoising strategies with ease.

Similar functionality provided by the `load_confounds` and `load_confounds_strategy` APIs are included in other fMRIPrep-compatible fMRI processing software, such as C-PAC (Li et al., 2021), XCP-D (Adebimpe et al., 2023), and ENIGMA HALFpipe (Waller et al., 2022). Unlike our APIs, which focus on retrieving denoising regressors only, these softwares provide denoising utilities bundled in a full preprocessing workflow. The denoising regressor retrieval steps amongst those softwares are therefore not reusable and more difficult to reproduce. Our APIs provide the advantage that users can easily reuse the denoising strategies. In fact, XCP-D has adopted our APIs in their code base. A limitation of our APIs is that the implemented denoising strategies are limited to those covered by the regressors included in fMRIPrep. With the constant development of denoising strategies, what the APIs provide will always lag behind the advancement of the field. However, as a trade-off, we can ensure the quality and robustness of the implementation.

### Denoising strategy

In order to summarize our results, we created a table ranking strategies from best to worst, based on four benchmark metrics, across datasets and fMRIPrep versions (see Figure 12). The ranking of the loss of temporal degrees of freedom is an important consideration accompanying the remaining metrics, as any denoising strategy aims at a particular trade-off between the amount of noise removed and the preservation of degrees of freedom for signals. Aside from the loss of temporal degrees of freedom, the `baseline` strategy consistently performs the worst, as expected, with the notable exception of `aroma` performing worst on QC-FC.

**Figure 12.**
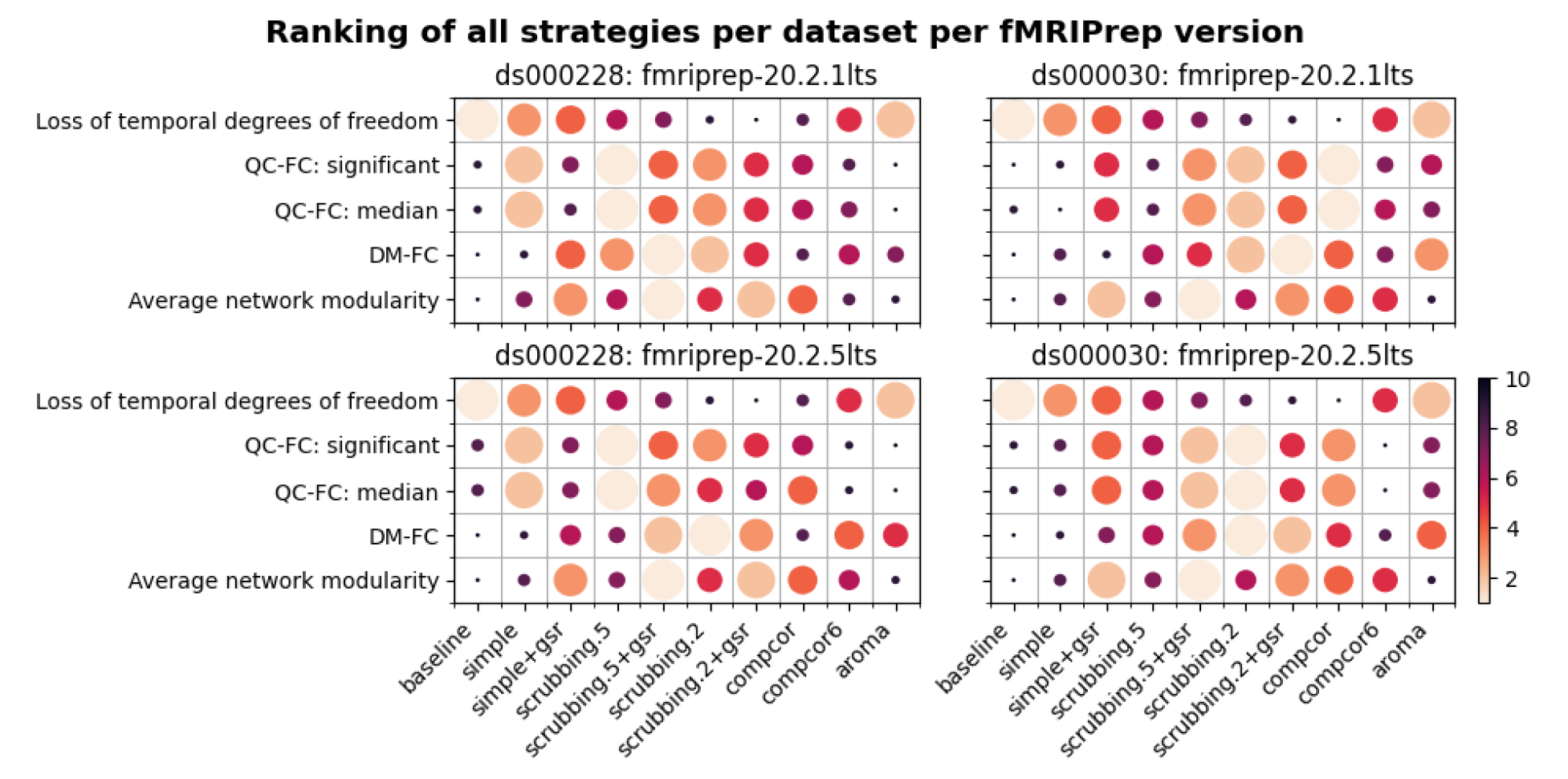
Ranking of all denoising strategies. *We ranked four metrics from best to worst. Larger circles with brighter color represent higher ranking. Metric “correlation between network modularity and motion” has been excluded from the summary as it is potentially a poor measure. Loss of temporal degrees of freedom is a crucial measure that should be taken into account alongside the metric rankings*.

The `simple+gsr` strategy is not the best for any particular individual evaluation metric, but it performed consistently well across metrics, datasets and software versions. The loss in degrees of freedom `simple` (26 + number of cosine terms) and `simple+gsr` (27+number of cosine terms) used slightly more regressors than `aroma`, and had markedly lesser loss than `scrubbing` methods. `simple+gsr` is consistently better than other data-driven strategies, which makes it the best choice for analysis that requires low loss of degrees of freedom and also preserve continuous sampling time series (which is broken by `scrubbing`).

Scrubbing based strategies are the best when it comes to minimizing the impact of motion, with a cost of higher loss in degrees of freedom. We found that scrubbing with an aggressive 0.2 mm threshold (`scrubbing.2`) mitigates distance dependency well consistently, regardless of the group of subjects. Despite excluding data with the same standard on both datasets, the child-dominant sample (*ds000228*) showed more volumes censored with the scrubbing strategy, and a liberal framewise displacement threshold showed sufficient ability to reduce the distance dependency of motion as observed in the original study of the strategy (Power et al., 2012). In a sample with higher motion, such as *ds000228*, a liberal scrubbing threshold reduced the impact of motion and performed similarly with a higher threshold. Taking the loss of degrees of freedom into consideration, we recommend a liberal scrubbing threshold rather than scrubbing with a stringent threshold for datasets with marked motion.

For the two anatomical CompCor strategies, `compcor` performs better than `compcor6`. The performance of `compcor6` is also not consistent across software versions in both datasets. However, `compcor` introduces large variability into the loss of degrees of freedom. In *ds000228*, the loss in temporal degrees of freedom is even higher than scrubbing with a stringent threshold. This result is consistent with the observation of Parkes and colleagues (2018) that anatomical CompCor is not sufficient for high motion data. Moreover, this observation puts one of the rationales in the original study, i.e., to reduce the loss in degrees of freedom, in question (Behzadi et al., 2007). In the absence of physiological recordings, our benchmark is not suitable to examine another property of CompCor, that is the ability to remove unwanted physiology signals (Behzadi et al., 2007). The datasets do not include physiology measures to perform alternative strategies such as RETROICOR to mitigate physiology signals explicitly.

In our results, `aroma` shows similar performance with the `simple` strategy across most metrics, with the exception of DM-FC (where it performs well). This strategy also featured a very low loss of degrees of freedom, making it a “gentle” denoising approach. Previous literature has recommended adding GSR as part of the ICA-AROMA regressors (Ciric et al., 2017; Parkes et al., 2018). An early version of this work did include a ICA-AROMA+GSR strategy, which performed very poorly (results not shown in the current version). This is a known consequence of implementation choices made in fMRIprep, which departs from the original recommended implementation of ICA-AROMA+GSR (Pruim, Mennes, van Rooij, et al., 2015)^6^. We strongly recommend fMRIPrep users to avoid fMRIPrep-generated GSR when using the ICA-AROMA strategy. It is also worth noting that fMRIPrep will drop the support for ICA-AROMA from version 23.1.0^7^.

Strategies including GSR produced connectomes with higher network modularity compared to their counterparts without GSR. There is no systematic trend of whether GSR improves the denoising strategies based on the remaining impact of motion. The result is consistent with the fact that global signal regression increases the number of negative connections in a functional connectome (see Nilearn examples visualizing connectomes with and without global signal regression^8^) by shifting the distribution of correlation coefficients to be approximately zero-centered (Murphy & Fox, 2017). A clear benefit of GSR is thus to emphasize the network structure, but its benefits for denoising can vary. Some strategies, such as `simple`, seem to benefit greatly from the addition of GSR.

### Re-executable research object

We created a re-executable denoising benchmark with two main outcomes. Firstly, we created a reusable code base that will ensure the robustness of the procedure. The current benchmark includes several parameters, from the choices of atlases, denoising strategies, fMRIPrep versions, to datasets. The code for connectome generation and denoising metric calculation is written as an installable Python library (https://github.com/SIMEXP/fmriprep-denoise-benchmark). Customized scripts to deploy the process for each combination of the parameters are also generated by reusable Python functions. The full workflow can be executed on the two benchmark datasets preprocessed by any release from the fMRIPrep LTS series. Full documentation to re-execute the workflow, from fetching datasets to running the analysis, is available as part of the research object^9^. Secondly, we created an interactive Jupyter Book (Granger & Perez, 2021) hosted on NeuroLibre (Karakuzu et al., 2022) for users to freely browse the results with finer details. All figures in this report can be rebuilt with the provided Makefile, handling data download and the report generation. Taken together, it is possible to reproduce the results of this manuscript, starting from raw data down to final figures, and update the entire manuscript on future releases of fMRIPrep, turning this research object into a living publication rather than a snapshot of current software destined for quick deprecation.

There are additional benefits from creating a re-executable denoising benchmark. The code for the current project is a good prototype of different BIDS-apps for post processing (Gorgolewski et al., 2017): a connectome generation BIDS-app and a denoising metric generation BIDS-app. BIDS-app is easier for user adoption under the BIDS convention and can expand the scope of the benchmark from the two datasets shown here to any BIDS-compliant dataset. The process of creating this benchmark also provides valuable first hand information about runtime, and the impact of atlas choice on computational costs, which we did not cover here but has big practical implications. High dimensional probabilistic atlases require four times more RAM than discrete segment atlases. For metric generation, high dimensional atlases can have a runtime up to 24 hours compared to 1 hour for low dimensional atlases. There is thus a very concrete “reproducibility” cost which comes with high-resolution and probabilistic atlases. The issue is rarely mentioned regarding the reproducibility of science, yet can be a real obstacle to actual reproduction. Future editions of the workflow will be built with runtime optimization in mind and potentially improve the code base for upstream projects, such as fMRIPrep.

### Continual evaluation of software versions

Our benchmark results on two versions of the long-term support (LTS) release of fMRIPrep reveals similar trends in the metrics, but some inconsistency. Between the two datasets, *ds000228* showed more consistent results than *ds000030* across two LTS releases (see Figure 12). The marked difference in *ds000030* was likely the result of a bug fix implemented in 20.2.2LTS^10,11^ and that *ds000030* had been reported as an affected dataset. The results from the data-driven strategies in both datasets demonstrated inconsistent relative difference when comparing to the `baseline` strategy. This piece of work is a new addition to the existing literature on the heterogeneity of results found through research software testing (Bowring et al., 2019; Gronenschild et al., 2012). Beyond mere numerical instabilities, we show that the qualitative conclusions of an evaluation benchmark do not necessarily generalize to different software packages or even versions of the same package. Our results thus highlight the importance of continuous evaluation of research software at each major step of its life cycle.

Rebuilding this paper on future fMRIPrep releases can be used to perform such continuous evaluation for future releases of fMRIprep. This benchmark is thus a hybrid contribution, being as much research paper as it is a software development tool. We still recommend several aspects of improvements to better achieve this goal for future similar efforts. Firstly the API will need to be kept up to date with fMRIPrep releases. The current code will be applicable for 20.2.x series up to September 2024. For fMRIPrep release beyond the LTS version, as long as the API in Nilearn is maintained, the code used to generate all current reports can be applied to the same two datasets. With the high number of tunable parameters (denoise strategies, atlases, software versions), a framework allowing parameter configuration, such as Hydra^12^, would help better manage and expand the benchmark. The current benchmark generates jobs through metadata stored in python dictionaries. By adapting a framework like Hydra, one can deploy the benchmark analysis with a simplified interface.

Finally, we note that all of the components necessary to implement our reproducible benchmarks are generic, i.e. software containers, data versioning, open source code with standard APIs, Jupyter Books and the Neurolibre preprint server. Beyond the particularities of fMRI denoising and the fMRIprep implementation, we thus believe that this work proposes an approach to implement reproducible benchmarks that is widely applicable, and would likely be beneficial in all scientific fields with heavy reliance on computational tools.

## Conclusions

This work introduces new software libraries to systematically evaluate the impact of a wide range of denoising strategies on resting state functional connectivity across datasets, and versions of the fMRIPrep preprocessing pipeline. We used this software infrastructure to implement a fully reproducible benchmark of denoising strategies on two datasets with varied characteristics, including age, motion level and the presence of clinical diagnoses. We would like to provide two strategy recommendations based on this benchmark, depending on a key consideration: whether preserving continuous sampling time series is needed (e.g. to train auto-regressive models) or not (e.g. to generate correlation coefficients across brain parcels). To preserve the continuous sampling property of time series, `simple+gsr` is the recommended strategy, especially for datasets with low motion, and appears to be robust across software versions. If continuous temporal sampling is not a priority, `scrubbing.5` was the best strategy for datasets with marked motion where denoising quality can be favored over loss of temporal degrees of freedom. The performance of `aroma` departed from the conclusions of previous denoising benchmark works and only performed well in one metric. The denoising benchmark also demonstrated differences in the performance of specific denoising strategies across multiple fMRIPrep versions. We hope that our benchmark provides useful guidelines for the community and demonstrates the importance of continuous evaluation of denoising methods. Our software infrastructure is broadly applicable and may be beneficial to implement reproducible benchmarks across different tools and research fields.

## Materials and Methods

### Datasets

Dataset *ds000228* (N = 155) contains fMRI scans of participants watching a silent version of a Pixar animated movie “Partly Cloudy”. The dataset includes 33 adult subjects (Age Mean(s.d.) = 24.8(5.3), range = 18 – 39; 20 female) and 122 child subjects (Age Mean(s.d.) = 6.7(2.3), range = 3.5 – 12.3; 64 female). T1w images were collected with the following parameters: TR = 2530 ms, TE = 1.64 ms, Flip Angle = 7°, 1 mm isotropic voxels. BOLD images were collected with the following parameters: TR = 2000 ms, TE = 30 ms, Flip Angle = 90°, 3 x 3 x 3.3 mm voxels. All images were acquired on a 3T Siemens Trio Tim Scanner. For more information on the dataset please refer to (Richardson et al., 2019).

Dataset *ds000030* includes multiple tasks collected from subjects with a variety of neuropsychiatric diagnosis, including ADHD, bipolar disorder, schizophrenia, and healthy controls. The current analysis focused on the resting-state scans only. Scans with an instrumental artifact (flagged under column ghost_NoGhost in participants.tsv) were excluded from the analysis pipeline. Of 272 subjects, 212 entered the preprocessing stage. Demographic information per condition can be found in *Table 2*. T1w images were collected with the following parameters: TR = 2530 ms, TE = 3.31 ms, Flip Angle = 7°, 1 mm isotropic voxels. BOLD images were collected with the following parameters: TR = 2000 ms, TE = 30 ms, Flip Angle = 90°, 3 x 3 x 4 mm voxels. All images were acquired on a 3T Siemens Trio Tim Scanner.

**Table 4.** Demographic information of ds000030.

|  | **Full sample** | **Healthy control** | **Schizophrenia** | **Bipolar disorder** | **ADHD** |
| --- | --- | --- | --- | --- | --- |
| N(female) | 212(98) | 106(54) | 30(8) | 41(19) | 35(17) |
| Age Mean(s.d.) | 33.2(9.3) | 31.8(8.9) | 37.2 (9.2) | 34.7 (8.9) | 32.5 (10.2) |
| Age Range | 21–50 | 21–50 | 22–49 | 21–50 | 21–50 |

### fMRI data preprocessing

We preprocessed fMRI data using fMRIPrep 20.2.1LTS and 20.2.5LTS through fMRIPrep-slurm (https://github.com/SIMEXP/fmriprep-slurm) with the following options:

~~~
--use-aroma \
--omp-nthreads 1 \
--nprocs 1 \
--random-seed 0 \
--output-spaces MNI152NLin2009cAsym MNI152NLin6Asym \
--output-layout bids \
--notrack \
--skip_bids_validation \
--write-graph \
--resource-monitor
~~~

For the full description generated by fMRIPrep, please see supplemental Jupyter Book^13^. We reported the primary outcomes using outputs from fMRIPrep 20.2.1LTS, and then investigated if the same conclusions can be observed in 20.2.5LTS.

### Choice of atlases

We extracted time series with regions of interest (ROI) defined by the following atlases: Gordon atlas (Gordon et al., 2016), Schaefer 7 network atlas (Schaefer et al., 2018), Multiresolution Intrinsic Segmentation Template (MIST) (Urchs et al., 2019) and Dictionary of Functional Modes (DiFuMo)(Dadi et al., 2020). All atlases were resampled to the resolution of the preprocessed functional data.

Since DiFuMo and MIST atlases can include networks with disjointed regions under the same label, we carried out further ROI extraction. Labels are presented with the original number of parcels. and we denote the number of extracted ROI in brackets. Gordon and Schaefer atlas parcels use isolated ROI, hence no further extraction was done. The Schaefer 1000 parcels atlas was excluded; regions were small enough that not all could be consistently resolved after resampling the atlas to the shape of the processed fMRI data.

- Gordon atlas: 333
- Schaefer atlas: 100, 200, 300, 400, 500, 600, 800
- MIST: 7, 12, 20, 36, 64, 122, 197, 325, 444, “ROI” (210 parcels, 122 split by the midline)
- DiFuMo atlas: 64 (114), 128 (200), 256 (372), 512 (637), 1024 (1158)

Processes involved here are implemented through Nilearn (Abraham et al., 2014). Time series were extracted using `nilearn.maskers.NiftiLabelsMasker` and `nilearn.maskers.NiftiMapsMasker`. Connectomes were calculated using Pearson’s Correlation, implemented through `nilearn.connectome.ConnectivityMeasure`.

### Participant exclusion based on motion

We performed data quality control to exclude subjects with excessive motion leading to unusable data. In the current report, we use framewise displacement as the metric to quantify motion. Framewise displacement indexes the movement of the head from one volume to the next. The movement includes the transitions on the three axes (*x*, *y*, *z*) and the respective rotation (α, β, γ). Rotational displacements are calculated as the displacement on the surface of a sphere of radius 50 mm (Power et al., 2012). fMRIPrep generates the framewise displacement based on the formula proposed in (Power et al., 2012). The framewise displacement, denoted as *FD*, at each time point *t* is expressed as: 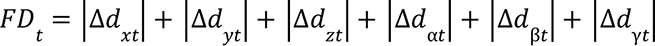 To ensure the analysis is performed in a realistic scenario we exclude subjects with high motion (Parkes et al., 2018) while retaining at least 1 minute of scan for functional connectome construction, defined by the following exclusion criteria: mean framewise displacement > 0.25 mm, above 80.0% of volumes removed while scrubbing with a 0.2 mm threshold.

### Confound regression strategies

Confound variables were retrieved using (i) a basic API that retrieves different classes of confound regressors, `nilearn.interfaces.fmriprep.load_confounds` (simplified as `load_confounds`); and (ii) a higher level wrapper to implement common strategies from the denoising literature, `nilearn.interfaces.fmriprep.load_confounds_strategy` (simplified as `load_confounds_strategy`). The following section describes the logic behind the design of the API. For documentation of the actual function, please see the latest version of Nilearn documentation (https://nilearn.github.io/stable/).

We evaluated common confound regression strategies that are possible through fMRIPrep-generated confound regressors, with the exception of ICA-AROMA + global signal regressor, as the implementation in fMRIPrep is not in line with the original implementation^14^. The connectome generated from high-pass filtered time series served as a baseline comparison. Confound variables were accessed using the API `load_confounds_strategy`. The detailed 10 strategies and a full breakdown of parameters used in these strategies is presented in Table 3.

### Evaluation of the outcome of denoising strategies

We first performed Pearson’s correlations to understand the overall numerical similarities of the denoised connectomes across different strategies. For each parcellation scheme, we computed a correlation matrix across the thirteen strategies. These correlation matrices were then averaged across the parcellation schemes within each dataset. The averaged correlation matrices were reordered into blocks of clusters with the function `scipy.cluster.hierarchy.linkage`. The aim was to provide an overview of the similarity of connectomes generated with the strategies.

We then used selected metrics described in the previous literature to evaluate the denoising results (Ciric et al., 2017; Parkes et al., 2018). After investigating the metrics with fMRIPrep version 20.2.1 LTS, we assessed whether the conclusions were consistent in 20.2.5 LTS.

### Loss in temporal degrees of freedom (Ciric et al., 2017; Yan et al., 2013)

The common analysis and denoising methods are based on linear regression. Using more nuisance regressors can capture additional sources of noise-related variance in the data and thus improve denoising. However, this comes at the expense of a loss of temporal degrees of freedom for statistical inference in further analysis. This may be an important point to consider alongside the denoising performance for researchers who wish to perform general linear model based analysis. Higher loss in temporal degrees of freedom can spuriously increase functional connectivity (Yan et al., 2013). Volume censoring-based and data-driven strategies (ICA-AROMA and some variations of CompCor) introduce variability to degrees of freedom and can bias group level comparisons (Ciric et al., 2017). We calculate the number of regressors used and number of censored volume loss. Depending on the length of the scan, the number of discrete cosine-basis regressors can differ. The number of discrete cosine-basis regressors will be denoted as *c* in the report (*c*_*ds*000228_ = 4, *c*_*ds*000030_ = 3). Simple, simple+gsr, compcor6 are the strategies with a fixed number of degrees of freedom loss. Scrubbing, compcor, aroma, and aroma+gsr strategies show variability depending on the number of noise components detected.

### Quality control / functional connectivity (QC-FC)

QC-FC (Power et al., 2015) quantifies the correlation between mean framewise displacement and functional connectivity. This is calculated by a partial correlation between mean framewise displacement and connectivity, with age and sex as covariates. The denoising methods should aim to reduce the QC-FC value. Significance tests associated with the partial correlations were performed, and correlations with P-values above the threshold of α = 0. 05 deemed significant. A version of this analysis corrected for multiple comparisons using the false discovery rate (Benjamini & Hochberg, 1995) is available in the Neurolibre preprint (Wang et al., 2023).

### Distance-dependent effects of motion on functional connectivity (DM-FC)

To determine the residual distance-dependence of subject movement, we first calculated the Euclidean distance between the centers of mass of each pair of parcels (Power et al., 2012). Closer parcels generally exhibit greater impact of motion on connectivity. We then correlated the distance separating each pair of parcels and the associated QC-FC correlation of the edge connecting those parcels. We report the absolute correlation values and expect to see a general trend toward zero correlation after confound regression.

### Network modularity

Confound regressors have the potential to remove real signals in addition to motion-related noise. In order to evaluate this possibility, we computed modularity quality, an explicit quantification of the degree to which there are structured subnetworks in a given network - in this case the denoised connectome (Satterthwaite et al., 2012). Modularity quality is quantified by graph community detection based on the Louvain method (Rubinov & Sporns, 2010), implemented in the Brain Connectivity Toolbox (Rubinov & Sporns, 2010). If confound regression and censoring were removing real signals in addition to motion-related noise, we would expect modularity to decline. To understand the extent of correlation between modularity and motion, we computed the partial correlation between subjects’ modularity values and mean framewise displacement, with age and sex as covariates.

### Contributions

The initial API was started by Hanad Sharmarke and Pierre Bellec. The implementation was completed by Hao-Ting Wang, Steven Meisler, François Paugam, and Pierre Bellec. Hao-Ting Wang migrated the code base to Nilearn. Nicolas Gensollen and Bertrand Thirion reviewed the code migrated to Nilearn. We thank Chris Markiewicz for feedback related to fMRIPrep.

Hao-Ting Wang and Pierre Bellec drafted the initial version of the paper, with critical feedback from Natasha Clarke. All co-authors contributed to and approved the final version of the manuscript.

Please see the original repository (https://github.com/SIMEXP/load_confounds#contributors-) for a history of initial development and contributors, and this issue (https://github.com/nilearn/nilearn/issues/2777) for a history of the integration in Nilearn and all the linked Pull Requests.

## Data and Code availability statements

Research code is available on GitHub repository (https://github.com/SIMEXP/fmriprep-denoise-benchmark). Datasets used in the current study are existing open access datasets on OpenNeuro (https://openneuro.org/datasets/ds000228/versions/1.1.0, https://openneuro.org/datasets/ds000030/versions/1.0.0). All metadata and summary statistics are available on Zenodo (https://doi.org/10.5281/zenodo.6941757). Retrieval of the data mentioned above are retrievable through the code repository and the Neurolibre preprint (https://doi.org/10.55458/neurolibre.00012).

## Ethics Statements

N/A

## Acknowledgements

The project is funded by IVADO PRF3, CCNA and Courtois Foundation, the neuromind collaboration. HTW and NC funded by Institut de valorisation des données (IVADO) postdoctoral research funding. SLM was funded by the National Institute on Deafness and Other Communication Disorders (NIDCD; Grant 5T32DC000038). CJM funded by NIMH 5R24MH117179. FP funded by Courtois Neuromod. PB funded by Fonds de Recherche du Québec - Santé (FRQ-S).

## Supplemental Materials

### Annex A. Common families of confound regressors

Mitigating the variance introduced by confounding fluctuations is necessary to gain meaningful measures of brain connectivity (Power et al., 2015). The most common method to minimize the impact of confounds is the use of linear regression (Friston et al., 1994) to regress out nuisance signals. These nuisance signals fall into one of three general categories commonly acknowledged in the literature: head motion, physiological noise (cardiac and respiratory), and instrumental noises from the MRI. The most common confound regressors are extracted following basic processing steps (i.e., motion correction, field unwarping, normalization, bias field correction, and brain extraction):

- Motion realignment measures capture head motion, a well-known source of disturbances in fMRI signals (Friston et al., 1996; Hajnal et al., 1994) which causes distance-dependent signal correlations and introduces systematic bias in group comparisons (Power et al., 2012; Satterthwaite et al., 2012; Van Dijk et al., 2012). Six rigid-body motion parameters (3 translations and 3 rotations) are typically estimated relative to a reference image and used as confound regressors (Friston et al., 1996).
- Non-grey matter tissue signals (such as *white matter* and *cerebrospinal fluid*) are unlikely to reflect neuronal activity and can be dominated by a mixture of motion and physiological artifacts (Fox et al., 2005). This type of signal is captured by averaging signals within anatomically-derived masks.
- The global signal is a confound regressor extracted from averaging signals within the *full brain* volume (Fox et al., 2005). The global signal clearly captures motion and physiological fluctuations, but is also sensitive to global neural activity, making it a controversial choice for regression (Power et al., 2017; Saad et al., 2012).
- Scrubbing (Power et al., 2012) is a volume censoring approach to remove high motion segments in which the framewise displacement (see Materials and Methods section Participant exclusion based on motion) exceeds some threshold. The scrubbing approach is applied alongside head motion parameters and tissue signal regressors.
- Temporal high-pass filtering accounts for low-frequency signal drifts introduced by *physiological and scanner noise sources*.

The family of motion, non-grey matter and global signal regressors can be further expanded using their first temporal derivatives and their quadratic (square) terms (Satterthwaite et al., 2013) to capture potential non-linear effects of these noise sources. Optimal denoising results often require full expansion of head motion parameters (both derivatives and squares).

Aside from regressors directly modeling noise derived from realignment measures or anatomical properties, other approaches capture the impact of motion and non-neuronal physiological activity through data-driven methods:

- The principal component-based method CompCor (Behzadi et al., 2007; Muschelli et al., 2014) extracts reduced components from white matter and cerebrospinal fluid masks to estimate non-neuronal activity.
- Independent component analysis-based methods estimate spatially independent components representing brain activity and noise. To identify the components related to head motion, researchers used a data-driven classifier (ICA-FIX (Salimi-Khorshidi et al., 2014)) or a pre-trained model (ICA-AROMA (Pruim, Mennes, van Rooij, et al., 2015)).

Despite the abundance of measures for non-neural signals, there is no one class of measures that can capture all known noise. A combination of nuisance regressors is thus needed to address the different sources of noise, and different common strategies have been put forth in the literature.

### Annex B. Common denoising strategies

Researchers developed and proposed different strategies with the aim of improving denoising approaches. A denoising strategy typically involves the selection of a subset of nuisance regressors from the broad list presented above. Head motion combined with non-grey matter tissue signals is one of the most basic approaches (Fox et al., 2005). Scrubbing combines the basic approach above with volume censoring, which can be implemented either by removing time points entirely (Power et al., 2012) or by adding “spike” nuisance regressors (Lemieux et al., 2007; Siegel et al., 2014). Anatomical CompCor regressors are usually applied along with the basic head motion parameters (Muschelli et al., 2014). ICA-AROMA requires two steps of denoising: (i) a partial regression to remove variance associated with noise ICA-AROMA components inside a full ICA decomposition preceded by (ii) a linear regression including the basic average of white matter and the cerebrospinal fluid signals regressors (Pruim, Mennes, van Rooij, et al., 2015). As there exists multiple variants of most categories of confounds, e.g. degree of expansion in motion parameters, a large number of specific variants do exist for each strategy through combinatorial effects. To provide objective guidance to practitioners in their selection of an effective denoising strategy, comprehensive denoising benchmarks have emerged in the functional connectivity research literature.

### Annex C. Evaluation of denoising strategies

Denoising benchmarks use several metrics to evaluate the effectiveness of denoising strategies on functional connectivity, commonly including loss of temporal degrees of freedom, residual motion effects, and impact on network properties, such as modularity. The ability of different strategies to remove confounds is heterogeneous (Ciric et al., 2017), and no single strategy dominates across all evaluation metrics. Several benchmarks still recommended ICA-AROMA due limited loss of degrees of freedom compared to scrubbing-based methods (Parkes et al., 2018), possibly in combination with global signal regression for maximal reduction of motion artifacts (Burgess et al., 2016; Ciric et al., 2017). However, a denoising strategy can perform differently due to factors that strongly correlate with motion, such as psychiatric conditions, age, or the choice of subsequent analytical technique. For example, CompCor may only be maximally effective in low-motion data (Behzadi et al., 2007). Volume-censoring-based strategies are unsuitable for time series analysis where a uniform sampling of signals is required, such as most time-frequency analysis implementations. Researchers thus need to evaluate the best denoising strategy based on the available benchmarks, the profile of their data and their choice of analytical techniques.

The existing benchmarks have a few pitfalls that limit the integration to each researcher’s unique needs. The benchmark research and denoising methods development are conducted on in-house preprocessing solutions with different datasets. From the research standpoint, this is not necessarily a pitfall as it showcases that workflows built upon different software following the general shared principle for preprocessing have converging conclusions. However, this is a problem for user adoption of the recommended strategy to their choice of preprocessing workflow. To correctly implement the best approach for their study, researchers need to understand the extensive literature and then construct the workflow. Another pitfall is that results in benchmarks are subject to the scope of datasets evaluated. As researchers use in-house tools and analysis code, there is no direct way to apply the suggested strategy or generate the benchmark statistics for evaluation on a new dataset. The denoising benchmark literature provides a good overview of the progress of methods in the field of resting state functional connectivity, but falls short in providing a reproducible path for the general community to adapt the results to modern solutions of preprocessing, such as fMRIPrep.

## Footnotes

1 Nilearn documentation for `load_confounds`: https://nilearn.github.io/stable/modules/generated/nilearn.interfaces.fmriprep.load_confounds_strategy.html#nilearn.interfaces.fmriprep.load_confounds

2 Nilearn documentation for `load_confounds_strategy`: https://nilearn.github.io/stable/modules/generated/nilearn.interfaces.fmriprep.load_confounds_strategy.html#nilearn.interfaces.fmriprep.load_confounds_strategy

3 https://preprint.neurolibre.org/10.55458/neurolibre.00012/

4 Move `sample_mask` to transform method in maskers, handle `sample_mask` in `signal.clean` https://github.com/nilearn/nilearn/pull/2858

5 https://nilearn.github.io/stable/auto_examples/03_connectivity/plot_signal_extraction.html#sphx-glr-auto-examples-03-connectivity-plot-signal-extraction-py

6 In fMRIPrep, the whole brain global signal regressor and the estimated head-motion parameters were calculated on the output from their regular pipeline (i.e., before denoising), which is inconsistent with the original proposal (Pruim, Mennes, van Rooij, et al., 2015) and not suitable for combining with ICA-AROMA. Based on the original implementation, the GSR should be calculated from the time series after removing the variance of ICA-AROMA regressors (Pruim, Mennes, van Rooij, et al., 2015). Otherwise,There is a possibility that the global signal regressor reintroduced motion to the data (Lindquist et al., 2019).

7 https://github.com/nipreps/fmriprep/issues/2936

8 https://nilearn.github.io/stable/auto_examples/03_connectivity/plot_signal_extraction.html#the-impact-of-global-signal-removal

9 https://preprint.neurolibre.org/10.55458/neurolibre.00012/docs/tldr.html

10 See: https://github.com/nipreps/fmriprep/issues/2307

11 See #2444 in change log https://fmriprep.org/en/stable/changes.html#july-16-2021

12 https://github.com/facebookresearch/hydra

13 https://preprint.neurolibre.org/10.55458/neurolibre.00012/notebooks/CITATION.html

14 See ICA-AROMA related warning in https://fmriprep.org/en/20.2.1/outputs.html#confounds

